# FAN, the homolog of mammalian Apoptosis Antagonizing Transcription Factor AATF/Che-1 protein, is involved in safeguarding genome stability through the ATR induced pathway in *Arabidopsis*

**DOI:** 10.1101/2023.03.10.531693

**Authors:** Fang Liu, Bingshan Wang, Xiangyang Wang, Daofeng Dong, Lieven De Veylder, Shengdong Qi, Beatrix M. Horvath, Klaus Palme, Xugang Li

**Author notes:** Authors for correspondence: Xugang Li, Klaus Palme E-Mail: Xugang Li; Klaus Palme. These authors contributed equally to this work (Fang Liu, Bingshan Wang).

## Abstract

Cellular DNA can be damaged by endogenous or exogenous genotoxins. In plants, reduced genome stability can have a detrimental effect on development. Here, we show the identification of the *fan* mutant from an ethyl-methanesulfonate (EMS) mutagenized *Arabidopsis* Col-0 population on the basis of its short root and small leaf phenotype. The causative mutation was identified as a G-to-A transition at the border of the eighth intron and ninth exon of the *At5G61330* gene, resulting in a mis-spliced mRNA transcript. FAN is a homolog of the mammalian AATF/Che-1 protein consisting of conserved AATF/Che-1 and TRAUB domains in *Arabidopsis*. In the *fan* mutant, under normal conditions, we detected DNA damage and cell death response at the root tip, while hypersensitivity to the exogenously applied hydroxyurea (HU) compared to Col-0, suggesting that FAN plays a role in the DNA damage response (DDR). Furthermore, our results showed that FAN is involved in DDR pathway regulated by ATM/RAD53-RELATED (ATR). Taken together, these suggest that FAN is required for meristem maintenance and the DNA damage response.

## INTRODUCTION

In higher plants, new organs are generated from stem cells located in the root and shoot meristems. Within the root apical meristem (RAM), stem cells surround the rarely dividing quiescent centre (QC) cells, which express the *WUSCHEL-RELATED HOMEOBOX5* (*WOX5*) transcription factor gene (Haecker et al., 2004; Sarkar et al., 2007). Maintenance of the stem cell niche is regulated by several transcription factors. These include the AP2/PLT transcription factors, which provide an apical-basal patterning signal and are regulated by auxin, forming a gradient of *PLT* expression (Aida et al., 2004). On the other hand, the GRAS transcription factors SHORTROOT (SHR) and SCARECROW (SCR) provide a radial patterning signal (Di Laurenzio et al., 1996; Helariutta et al., 2000). In addition to these important transcription factors, genome integrity in stem cells and their progeny cells is required for meristem maintenance. For example, MAIN and MAIN Like family members are involved in maintaining genomic stability, and mutants exhibit short primary roots, disorganized RAM, and accumulated DNA breaks (Ühlken et al., 2014; Wenig et al., 2013). AtMMS21, a subunit of the STRUCTURAL MAINTENANCE OF CHROMOSOME5/6 complex, is involved in ameliorating DNA double strand breaks (DSBs) and maintaining the stem cell niche during root development (Xu et al., 2013). The *m56-1fas2-4* triple mutant lacking the H3/H4 histone chaperon CHROMATIN ASSEMBLY FACTOR-1 (CAF-1) and the H2A/H2B histone chaperone NAP1-RELATED PROTEIN1/2 (NRP1/2), which function synergistically in chromatin maintenance and genome maintenance, exhibited programmed cell death and severe short-root phenotype (Ma et al., 2018).

Like all living organisms on the Earth, plants suffer various endogenous and exogenous stresses, such as ultraviolet radiation, chemical mutagens, and reactive oxygen species, which threaten the integrity of their genomes (Roldan-Arjona and Ariza, 2009; Ryu et al., 2019). To prevent the transmission of damaged DNA to daughter cells, the ATM/RAD53-RELATED (ATR) and ATAXIA-TELANGIECTASIA MUTATED (ATM) kinases are activated in response to replication defects and DSBs, respectively (Culligan et al., 2004; Garcia et al., 2003), to ultimately induce DNA damage repair, programmed cell death, or endoreduplication. Different from animals, there is no orthologue of Chk1, Chk2 or CDC25, which are downstream factors of ATM or ATR, but the p53 functional homologue SUPPRESSOR OF GAMMA RESPONSE 1 (SOG1) and the kinase WEE1 exist in plants. SOG1 is phosphorylated by ATM in response to DNA damage (De Schutter et al., 2007; Yoshiyama et al., 2013), with the extent of Ser-Gln (SQ) phosphorylation in SOG1 mediating the strength of DNA damage responses (Yoshiyama et al., 2017). As a transcription factor, SOG1 transcriptionally regulates hundreds of genes that are involved in cell cycle regulation, endocycle, cell death and DNA repair. Meanwhile, the cell cycle regulatory kinase WEE1 controls cell cycle progression via the WEE1-RPL1-CYCDs and the WEE1-FBL17/SKP2-CKI-CDKs axis in response to replication stress (Pan et al., 2021; Wang et al., 2021). In addition, recent studies showed that RETINOBLASTOMA RELATED (RBR), the Arabidopsis Rb homolog, in addition to its well-known cell cycle control and the stem cell maintenance functions, also maintains genome integrity of root meristematic cells by mediating the localization of the repair protein RAD51 to DNA lesions and accumulating together with E2F, and possibly AtBRCA1 to γH2AX foci in *Arabidopsis* (Biedermann et al., 2017; Horvath et al., 2017). In addition, DSBs induce endoreplication through sets of cell cycle regulators regulated by ATM-SOG1 and ATR-SOG1 pathways (Adachi et al., 2011), and the selective killing of root and stem cells and their early descendants induced by zeocin is also triggered by ATM and ATR kinase (Fulcher and Sablowski, 2009).

Human AATF/Che-1 was first identified as a subunit of RNA polymerase II, that interacts with retinoblastoma (Rb) to repress its function (Fanciulli et al., 2000). In response to DNA damage, Che-1 is stabilized by phosphorylation by ATM/ATR and Chk2 (Bruno et al., 2006). Phosphorylated Che-1 activates p53 and p21 transcription to maintain the G2/M checkpoint (Bruno et al., 2006). In addition to p53 transcriptional regulation, as a cofactor, phosphorylated Che-1 binds to p53 and forms a ternary complex with Brca1 in the first hours of DNA damage, resulting in transcription of growth arrest genes, and when cells accumulate excessive damaged DNA, p53 is released from the Che-1/p53/Brca1 complex to promote transcription of pro-apoptosis genes (Desantis et al., 2015). However, in tumor cells, mutant p53 (mtp53) inhibits the transcriptionally active forms of the p53 homologs p73 and p63 and phosphorylated Che-1 promotes transcription of mutant p53, resulting in tumor cell survival (Bruno et al., 2010). This anti-apoptosis function in tumor cells makes Che-1 a theoretical therapeutic target for cancer treatment. In general, AATF/Che-1 plays important roles in the regulation of proliferation and survival in the DNA damage response pathway in mammals. In addition to its role in the DNA damage response pathway, AATF/Che-1 has been reported to be a key component of ribosome biogenesis. Recently, researchers discovered that AATF/Che-1 interacts with NGDN and NOL10 to form a complex called the ANN complex. This nucleolar complex is involved in the synthesis of 40S ribosomal subunits (Bammert et al., 2016). Also, AATF/Che-1 interacts with several RNA species and proteins such as 45S pre-rRNA, snoRNAs, ribosome biogenesis mRNAs, rRNA processing proteins and ribosomal proteins, that are important for ribosome biogenesis (Kaiser et al., 2019). In addition, Che-1 affects rDNA transcription by binding to the RNA polymerase Ⅰ machinery (Sorino et al., 2020).

Although AATF/Che-1 has been well studied in animals, little is known about its plant homologue. Here, we describe the isolation of the mutant, named *fan*, which exhibits severe aberrations in root architecture, root growth inhibition and severe defects and cell death in the root meristem. Based on the phenotypic changes, we propose that FAN, the *Arabidopsis* AATF/Che-1 homolog, is involved in root development and DNA damage response and is localized in the nucleolus. Our analysis of *atr-2;fan* double mutant shows that FAN is responsible for genome integrity through the ATR pathway.

## RESULTS

### The *fan* mutant has a short primary root and a disorganized stem cell niche (SCN)

The mutant, named *fan*, which we isolated from the EMS mutagenized *Arabidopsis* Col-0 population, showed retarded growth, including shorter primary roots and smaller cotyledons compared to Col-0 (Figure 1A). To eliminate background mutations, we backcrossed the homozygous *fan* mutant with Col-0. By assessing the F1 phenotype and the segregation rate of the *fan* phenotype of 3:1 (221:68, Chi-squared test, p=0.56), we found that *fan* is a recessive allele. Compared to the control plants, the root meristem length of the backcrossed *fan* mutant was already shorter at 4 DAG, and decreased further with time (Figure 1B, C). The number of transit amplifying cells in the cortex layer was significantly (*P*-value < 0.01) reduced in the *fan* mutant compared to Col-0 at 4 DAG. Although the number of cells slightly increased in Col-0 at 6 DAG, it continued to decrease in *fan* (Figure 1B, D), indicating progressive consumption of the meristem. Following root growth for 8 days after germination (DAG), we showed that the primary root length of *fan* was significantly shorter than that of Col-0 (Figure 1E). These data suggest that the transit amplifying cells in *fan* differentiate at an earlier stage of development.

**Figure 1.**
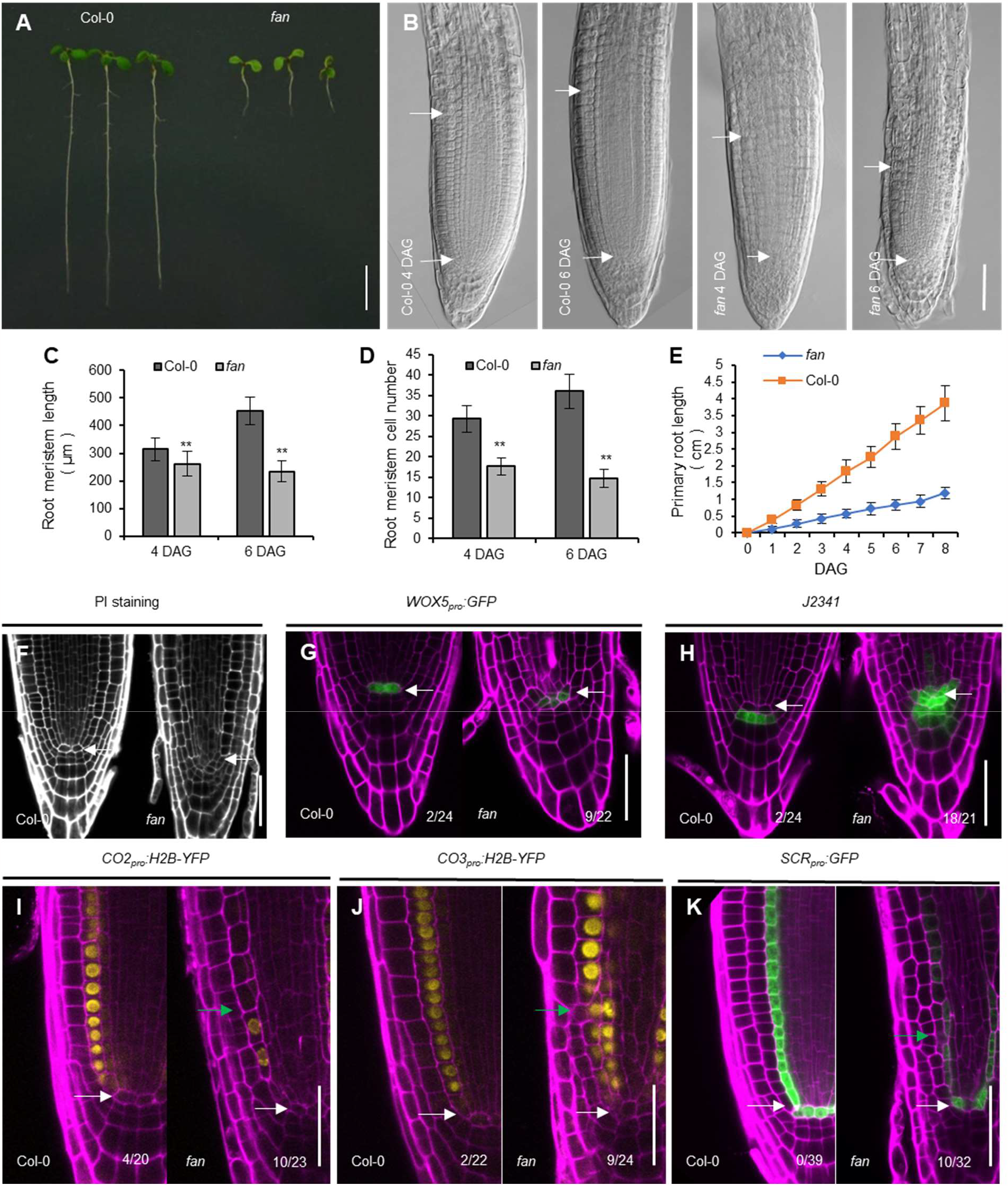
Primary root growth and stem cell niche (SCN) maintenance are inhibited in the *fan* mutant. (**A**) Growth habit of the *fan* mutant compared to the control, Col-0 at 6 DAG under normal growth conditions. Scale bar = 1 cm. (**B**) Representative images of differential interference contrast (DIC) microscopy showing the root meristem of Col-0 and *fan* (4 and 6 DAG). Scale bar = 50 µm; arrowheads point to QC and first elongated cortical cell position. (**C**) and (**D**) Meristem length and number of amplifying cells in the cortex of Col-0 and *fan* (4 and 6 DAG). Data represent the mean with ± SD, n>20. Double asterisks indicate highly significant differences (*P*<0.01) between *fan* and Col-0 analyzed by Student’s *t* tests. (**E**) The graph shows the root growth of Col-0 and *fan* between 0-8 days after germination. Data represent the mean with ± SD; n=30. (**F**) to (**H**) Representative confocal images of propidium iodide (PI)-stained Col-0 and *fan* mutant root tips (**F**) PI only, (**G**) showing expression of *WOX5*_*pro*_*:GFP* while (**H**) of columella stem cell marker *J2341*. Scale bars = 50 µm, arrowheads point to QC. (**I**) to (**K**) Col-0 and *fan* roots expressing *CO2*_*pro*_*:H2B-YFP* (**I**), *CO3*_*pro*_*:H2B-YFP* (**J**), and *SCR*_*pro*_*:GFP*. The numbers show the number of seedlings that display abnormal expression patterns compared to the total number of examined seedlings. White arrowheads indicate to QC and green arrowheads point to abnormal cell division. Scale bars = 50 µm. Magenta, PI staining; yellow, YFP and green, GFP.

To investigate whether alterations in the SCN, specifically in the QC, lead to phenotypic changes during root meristem development, we stained the root meristem with propidium iodide (Figure 1F) and analyzed the expression of fluorescent stem cell markers (Figure 1G, H). An abnormal QC organization in the *fan* was clearly detected by PI staining (Figure 1F) and confirmed by the altered expression domain of the *WOX5*_*pro*_*:GFP* marker (Figure 1G). This observation was confirmed by the expression pattern of the independent QC-marker *QC25:GUS* (Figure S1).

To further investigate the defects of *fan* in the distal stem cell niche, we followed the *J2341* enhancer trap line, which in Col-0 shows expression in the columella initials. In the *fan* mutant, the expression domain of the columella stem cell marker *J2341* expanded into additional cells surrounding the QC cells (Figure 1H). To investigate the identity of these cells, we followed the expression of *CO2*_*pro*_*:H2B-YFP, CO3*_*pro*_*:H2B-YFP* and *SCR*_*pro*_*:GFP*, which are highly expressed in the cortex and endodermis (ten Hove et al., 2010). In the *fan* roots, all three markers showed abnormal localization and cell division patterns in the cortex and endodermis cell layers (Figure 1I, J, K). Taken together, these results suggest that FAN is required for the maintenance of the stem cell niche and root meristem.

### FAN encodes the mammalian homologue of AATF/Che-1 in Arabidopsis

The *FAN* gene was identified by map-based cloning. Preliminary mapping showed that the mutated site is located at the tail of chromosome 5 between markers MTE17 and MSN2 (Lukowitz et al., 2000). Using simple sequence length polymorphism (SSLP) markers and derived cleaved amplified polymorphic sequence (CAPS) markers, the mutation was fine-mapped, and all the genes in this region were sequenced and compared to the wild-type control. A mutated base pair was located in the gene annotated as At5g61330, at the boundary of its eighth intron and ninth exon, where the change from G to A resulted in RNA mis-splicing (Figure 2A). To assess the consequences of alternative splicing, we compared the sequences of the mutant and wild-type cDNAs by reverse transcription polymerase chain reaction (RT-PCR) using total RNA isolated from mutant and wild-type seedlings. Due to mis-splicing, the translation is prematurely terminated at 870 bp from the translational start site (Figure 2A), resulting in a protein being truncated by 290 amino acids.

**Figure 2.**
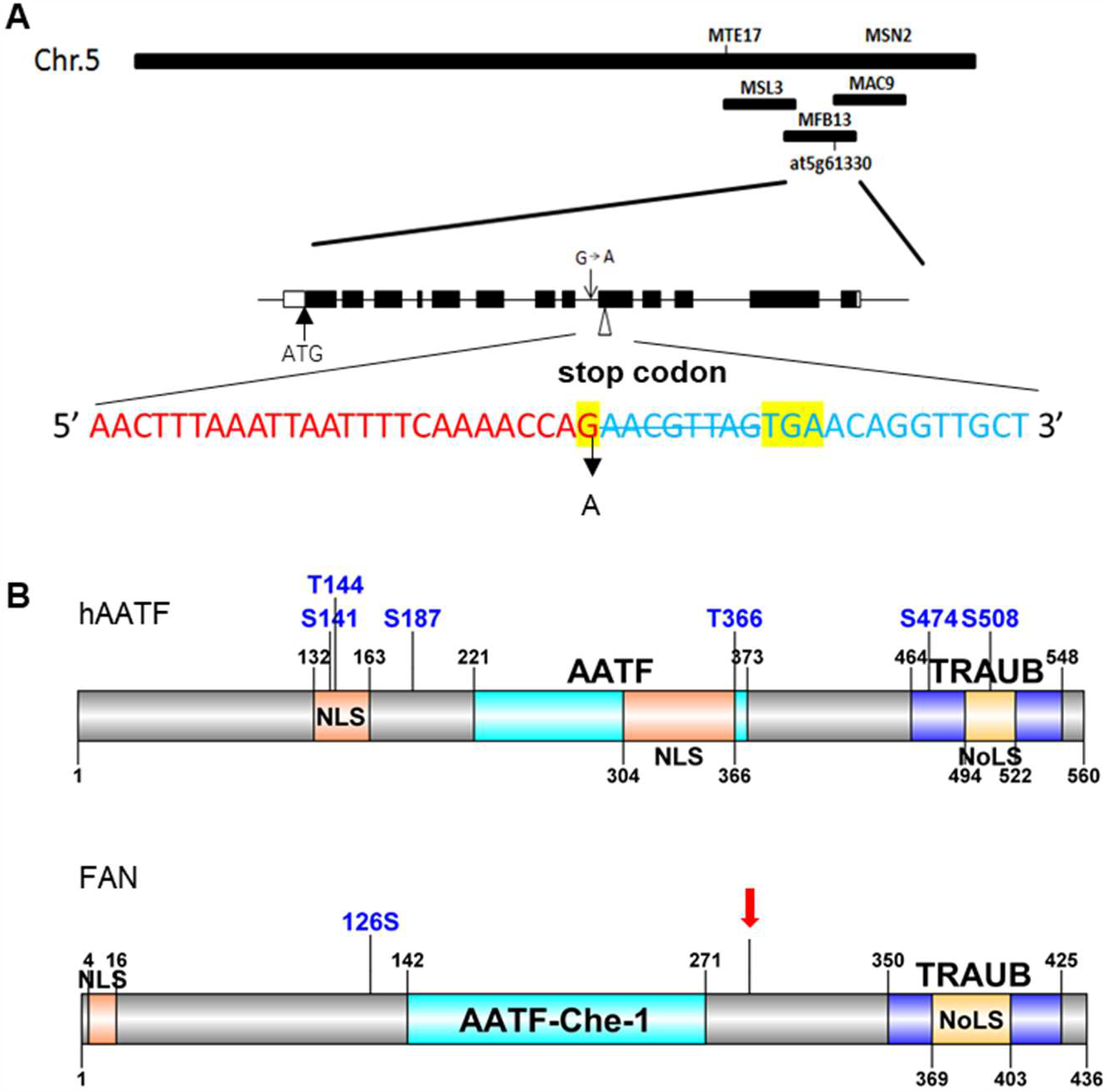
*FAN* encodes AATF/CHE-1 in *Arabidopsis*. (**A**) Upper section: description of markers. Middle section: intron-exon organization of *FAN*. Black boxes represent exons. The arrows indicate the position of the initiation codon and the mutated base pair site. Bottom section: Partial sequence of the eighth intron is shown in red and the ninth exon is shown in blue. By EMS mutagenesis, the last base of the eighth intron, G is mutated to A. This mutation leads to mis-splicing and with an in-frame stop codons TGA to a shorter transcript. AACGTTAG indicates the missing sequence ‘AACGTTAG’. The mutation site and stop codon are highlighted in yellow. (**B**) Comparison of the conserved protein boxes between the human AATF/Che-1 (hAATF) and FAN. Light blue box: AATF/Che-1 domain. Lila box: TRAUB. Orange box: Nuclear localization signal (NLS). Yellow box: Nucleolar localization sequence (NoLS). Red arrowhead: position of the point mutation in *FAN*. Blue letters and numbers indicate the confirmed and one of the potential phosphorylation sites on serine and threonine residues in hAATF and FAN respectively.

To confirm whether the mutation of the *FAN* gene is the cause of the observed growth defects in *fan* mutants, homozygous *fan* plants were transformed with the full-length genomic coding region of *FAN* expressed under its own promoter and using its 3’-UTR region (*FAN*_*pro*_*-FAN*_*g*_*-FAN*_*3’-UTR*_). Since the wild-type FAN protein fully complemented the short primary root and dwarf phenotype of *fan* (Figure S2), we could conclude that the mutation in the *FAN* gene was responsible for the mutant phenotype.

According to the public database (TAIR), *FAN* is predicted to encode an rRNA processing protein. Its coding sequence shares 41.7% sequence similarity with the human AATF/Che-1 protein, which encodes the Apoptosis Antagonizing Transcription Factor, with 26.5 % sequence identity (https://www.ebi.ac.uk/Tools/psa/emboss_needle/). FAN has two conserved protein domains, the N-terminal leucine-zipper region of AATF/Che-1 (at position 142 aa-271 aa) and the C-terminal TRAUB-domain (position 350 aa-425 aa) (Figure 2B). FAN is the only AATF/Che-1 in *Arabidopsis* that shows further conservation in both the animal and plant kingdoms (Figure S3).

### *FAN* is ubiquitously expressed and localized in the nucleolus

To study the expression of *FAN* in *Arabidopsis*, we generated *FAN*_*pro*_*:GUS* reporter lines and analyzed 3 independent transgenic lines. We observed GUS activity in young roots, young shoots, leaves, flowers, and siliques (Figure 3A-G). In particular, high expression was detected in the root apical meristem, consistent with the observed root phenotypes of the *fan* mutant. To investigate the cellular and subcellular localization of the protein, we generated translational reporter *FAN*_*pro*_*:FAN*_*g*_*-GFP* lines expressing the *FAN-GFP* fusion under its own promoter. Consistent with the GUS expression in the root, the FAN-GFP fusion protein showed high expression in the stem cells and the transit amplifying cells, but much weaker expression in the QC, columella cells, and the elongation zone, where cell division is less frequent (Figure 3H).

**Figure 3.**
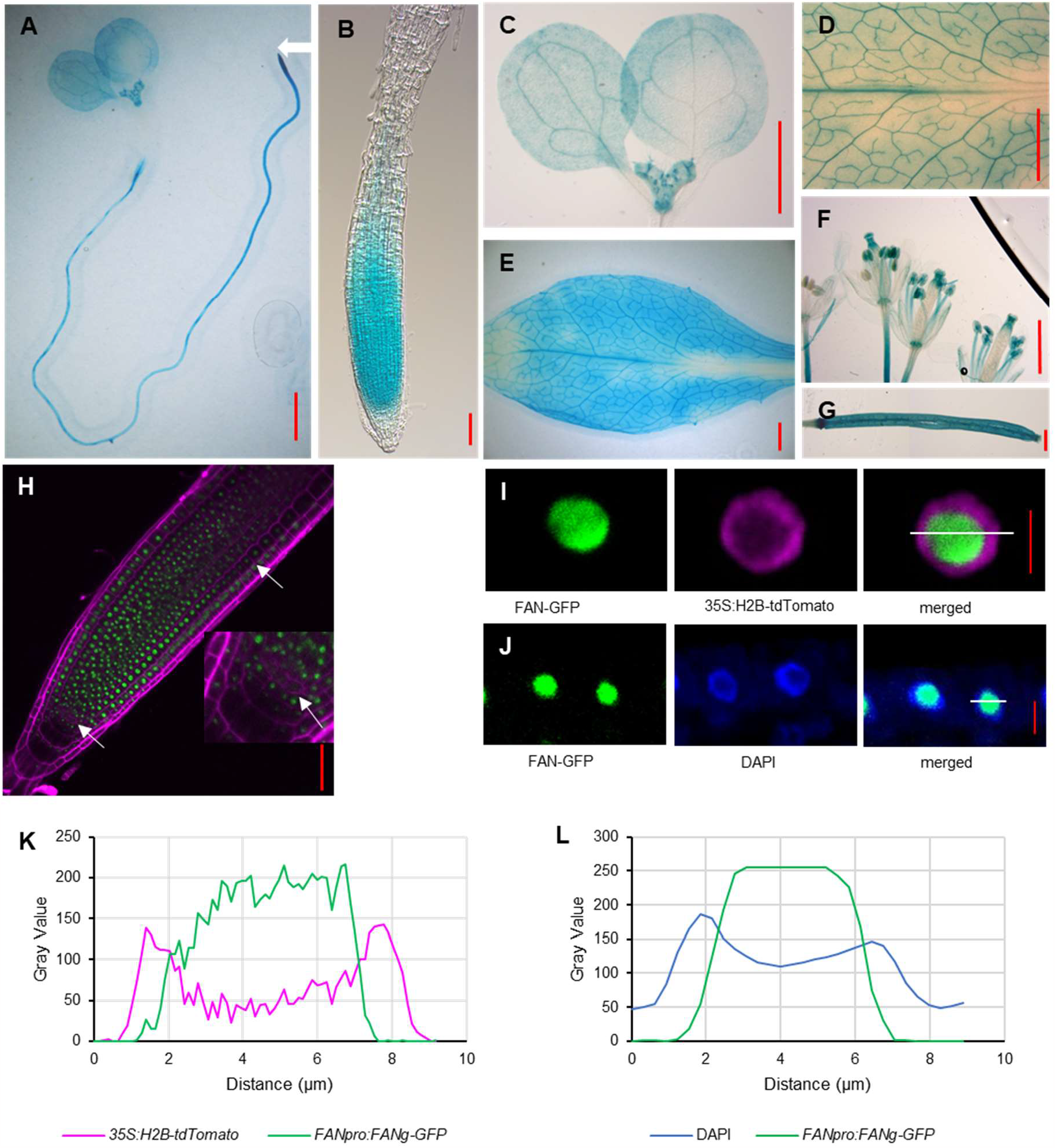
FAN is widely expressed and localized in nucleolus. (**A**) to (**G**) GUS staining was observed in (**A**) young seedling, (**B**) root, (**C**) young shoot, (**D**) and (**E**) mature rosette leaf and enlarged image, (**F**) flowers and (**G**) silique. The white arrowhead points to the root apical meristem. (**H**) The representative confocal image shows the expression of FAN in the root meristem of *FAN*_*pro*_*:FAN*_*g*_*-GFP* line (6 DAG), the enlarged section shows its QC localization. Magenta: PI staining; green, GFP fluorescence. Arrowheads point to QC and the first elongated cortex cell. Scale bars =1000 µm in (**A**), (**C**), (**D**), (**E**), (**F**) and (**G**), scale bars = 50 µm in (**B**) and (**H**). (**I**) Confocal images of the nuclear localization of the FAN-GFP in the *FAN*_*pro*_*:FAN*_*g*_*-GFP* line transformed with the *35s:H2B-tdTomato* marker. Magenta: H2B-tdTomato; green, FAN-GFP. Scale bar = 5 µm. (**J**) DAPI-stained nuclei of the *FAN*_*pro*_*:FAN*_*g*_*-GFP* line. Blue: DAPI; green, GFP. Scale bar = 5 µm. (**K**) and (**L**) Grey value distribution of the total number of pixels at the position of the white lines labelled in (**I**) and (**J**).

As shown in Figure 3H, the FAN-GFP fusion protein is localized to the nucleus. To further investigate the subcellular localization of FAN, we introgressed the nucleoplasm marker *35S:H2B-tdTomato* into the *FAN*_*pro*_*:FAN*_*g*_*-GFP* transgenic line. The labeled H2B allowed a more precise determination of the nuclear morphology, the nucleolus, and the surrounding nucleoplasm (Boisnard-Lorig et al., 2001). The confocal images of individual nuclei and their intensity profile indicated that the tdTomato and GFP signals did not overlap (Figure 3I, K). The fluorescent dye, 4’,6-diamidino-2-phenylindole (DAPI) visualizes nuclear DNA. Therefore, DAPI staining also shows the nucleolus region in the nucleus which appears as a dark area in the nucleus due to the low concentration of DNA in the nucleolus (Pontvianne et al., 2016). Our DAPI staining result confirmed that FAN is mainly localized in the nucleolus (Figure 3J, L). Taken together, our results demonstrate that FAN is highly localized to the nucleolus in dividing cells.

### Expression of key regulatory genes is altered in the *fan* roots

Because of the mutant root phenotype and the meristematic expression of FAN, we were interested in whether the alterations in root growth and development were accompanied by changes in the auxin/PLT pathway. To analyze the auxin distribution and levels in *fan*, we followed the auxin marker *DR5*_*rev*_*:GFP*. The auxin maximum in QC and the distribution in columella cells in *fan* did not differ from wild-type (Figure 4A). To follow the expression changes of the root master regulators, PLTs, we crossed the *PLT1*_*pro*_*:CFP* and *PLT2*_*pro*_*:CFP* marker lines into the *fan* mutant background. In the Col-0 background, the expression of *PLT1*_*pro*_*:CFP* and *PLT2*_*pro*_*:CFP* showed high expression in the stem cell niche, reached intermediate levels in rapidly dividing cells, and was low in elongated cells, which are required for maintenance of stem cell identity, promoting mitotic activity of stem cell daughters and cell differentiation (Galinha et al., 2007). As shown in Figure 4B and 4C, S4A, S4B, the expression level of *PLT1* and *PLT2* driven *CFP* expression was significantly reduced and restricted to the SCN in the *fan* mutant. Measuring the expression level of *PLT1* and *PLT2* by qRT-PCR in *fan* root tips confirmed our observation (Figure 4F). Similarly, the expression levels of the GRAS family genes *SCR* and *SHR* were also reduced compared to those in Col-0 (Figure 4D, 4E, and 4G; Supplementary Figure S4C and S4D). These results suggest that the aberrant root formation and SCN maintenance in the *fan* mutants is associated with the abnormal distribution of PLT1/PLT2, SCR, and SHR.

**Figure 4.**
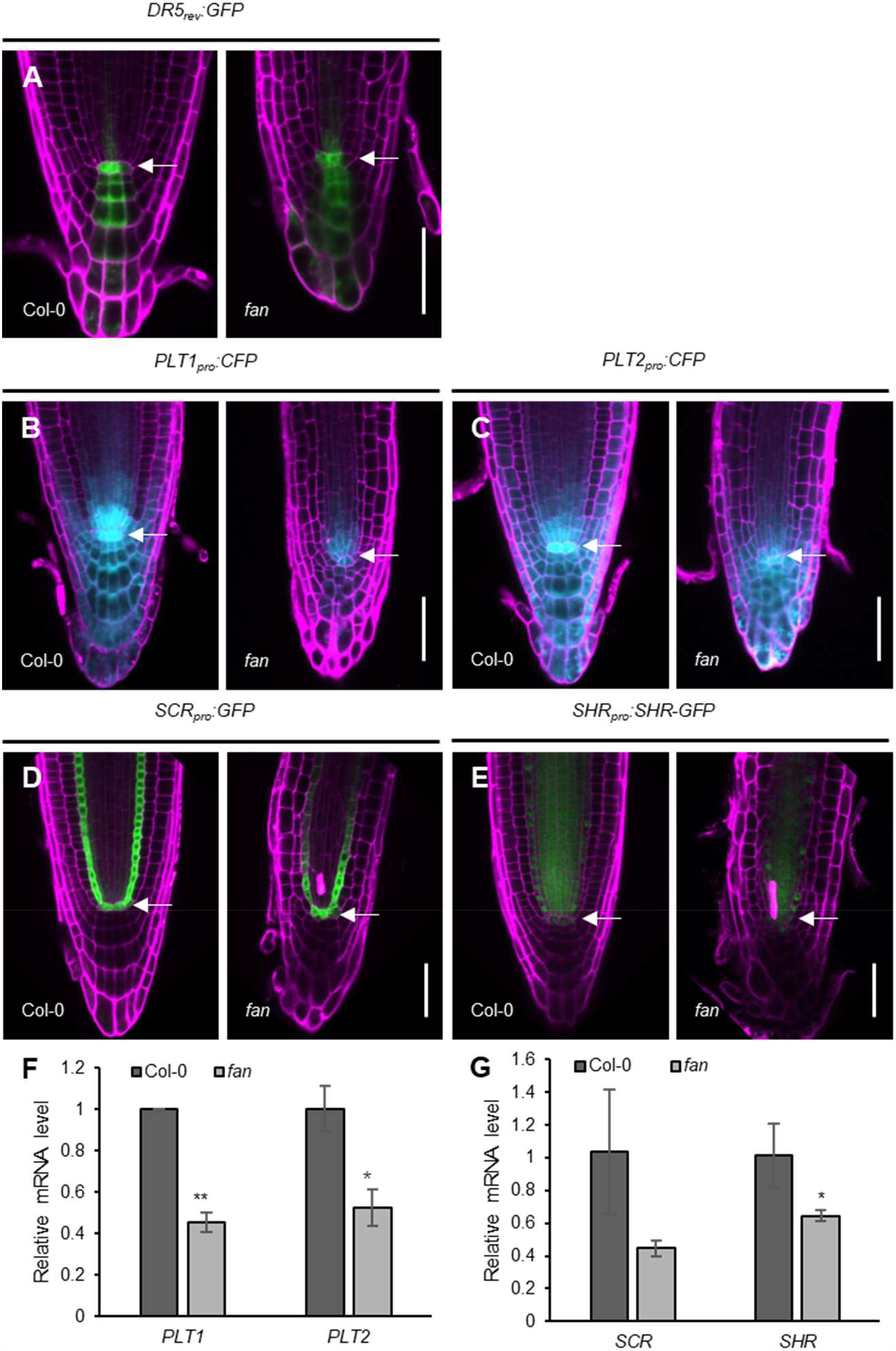
Auxin signaling and expression patterns of root developmental regulator genes are altered in *fan*. (**A**) Expression pattern of *DR5*_*rev*_*:GFP* in the root tip of Col-0 and *fan* (5 DAG). Scale bar = 50 µm; arrowheads point to QC. Magenta, PI staining; Green, GFP. (**B**) and (**C**) Expression pattern of *PLT1*_*pro*_*:CFP* (**B**) and *PLT2*_*pro*_*:CFP* (**C**) in the root tips of 5 DAG Col-0 and *fan*. Scale bars = 50 µm; arrowheads point to QC. Magenta, PI staining; Cyan, CFP. (**D**) and (**E**) Expression pattern of *SCR*_*pro*_*:GFP* (**D**) and *SHR*_*pro*_*:SHR-GFP* (**E**) in the root tips of 5 DAG Col-0 and *fan*. Scale bars = 50 µm; arrowheads point to QC. Magenta, PI staining; Green, GFP. (**F**) Relative mRNA levels of *PLT1* and *PLT2* measured by qRT-PCR in Col-0 and *fan*. 2 mm root tips were collected as samples. Data are mean ± SD of three biological replicates. Asterisks indicate significant differences compared with Col-0. (\*\**P* value < 0.01, \**P* value < 0.05, Student’s t test). (**G**) *SCR* and *SHR* relative mRNA levels measured by qRT-PCR in Col-0 and *fan*. 2 mm root tips were collected as samples. Data are mean ± SD of three biological replicates. Asterisks indicate significant difference compared to Col-0. (\**P* value < 0.05, Student’s t test).

### *fan* mutant show cell death and DNA damage and is hypersensitive to hydroxyurea

Similar to the mutations in the genes for *FAS1* (subunit of the counterpart of chromatin assembly factor-1) (Kaya et al., 2001) and *MMS21-1* (Xu et al., 2013), which are involved in stem cell niche and meristem maintenance and are critical for genome integrity, we also observed occasional cell death (42%) in the *fan* mutant’s meristem (Figure 5A).

**Figure 5.**
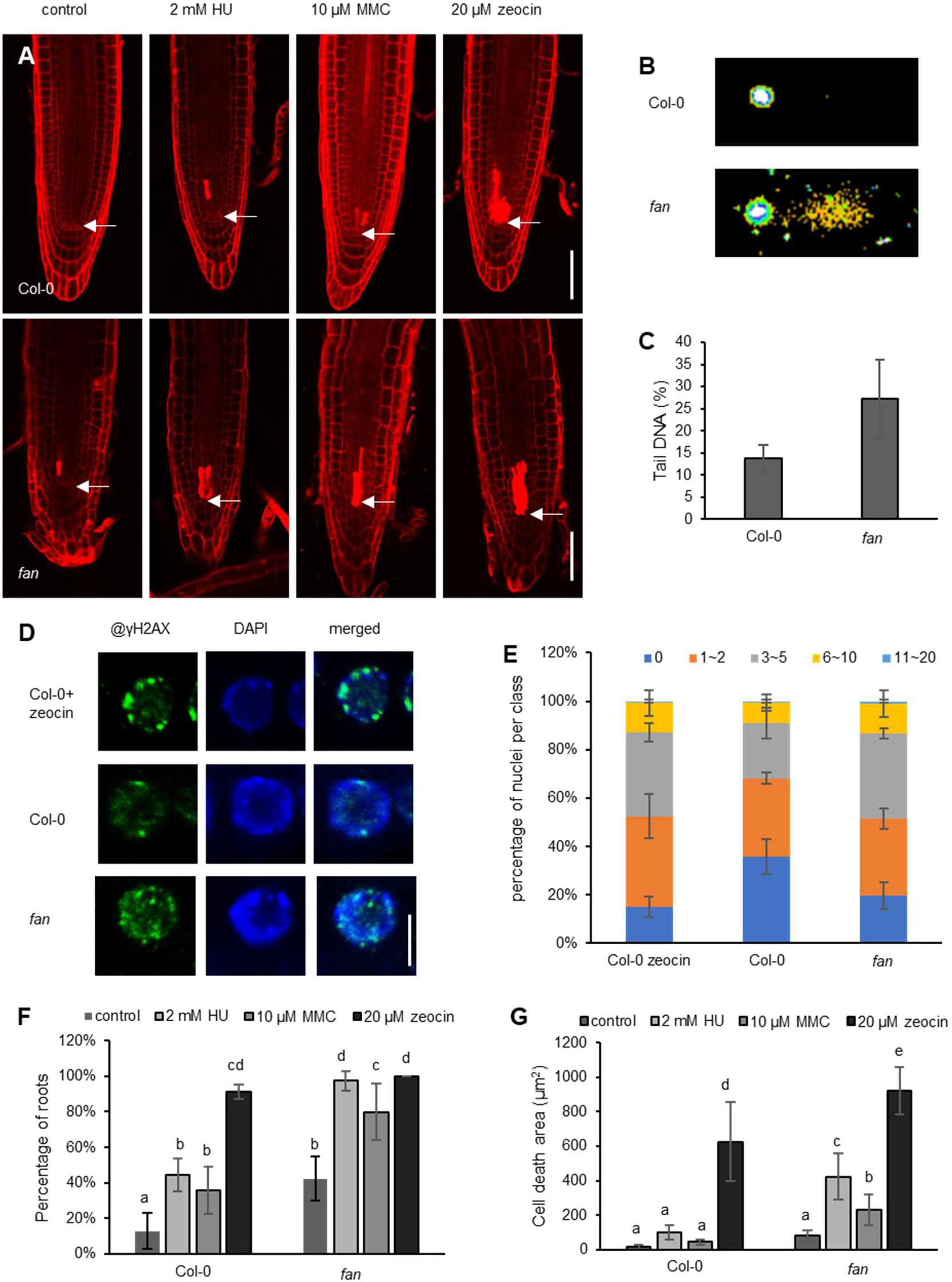
Increased cell death and broken DNA in *fan*. (**A**) Confocal images of PI-stained root tips of Col-0 and *fan* under normal growth conditions, treated with 2 mM HU treatment, 10 µM MMC and 20 µM zeocin. Seedlings (4 DAG) were transferred to 1/2 MS medium with or without DNA damaging agents. Scale bars = 50 µm; arrowheads point to QC position. (**B**) Representative comet assay images of nuclei from Col-0 and *fan* (7 DAG). (**C**) Levels of DNA fragments measured by the percentage of DNA in the tail of comets in the comet assay. Data are the means ± SD of three independent experiments. (**D**) γH2AX accumulation in the root tips of 5 DAG Col-0 on 1/2 MS medium containing 10 µM zeocin, Col-0 and *fan* grown under normal conditions measured by immunofluorescence staining. Blue, DAPI staining; green, γH2AX. Scale bar = 5 µm. (**E**) Quantification of γH2AX foci in Col-0 treated with 10 µM zeocin, Col-0 and *fan*. At least 100 nuclei were analyzed and grouped into 6 classes according to the number of γH2AX foci/nuclei. The data are from three independent experiments. Error bars indicate the SD. (**F**) and (**G**) The graphs show (**F**) the proportion of roots with cell death and (**G**) the mean area of dead cells after treatment compared to the control. Data are the mean ± SD from three independent experiments with at least 10 seedlings each. Columns with different letters are significantly different at *P*<0.05 (Duncan’s multiple range means comparisons).

To investigate whether the absence of FAN leads to genome instability, we first used the comet assay, and then followed the accumulation of the phosphorylated histone variant, γH2AX to detect DNA damage. The comet assay showed that the level of damaged DNA was higher in the *fan* mutant compared to the wild-type (Figure 5B, C). Immunofluorescence staining with the γH2AX antibody also showed that *fan* had increased DNA damage compared to the wild-type under normal growth conditions (Figure 5D, E). The number of γH2AX foci in the *fan* mutant was similar to that of Col-0 treated with 10 µM zeocin, a radiomimetic drug that causes DSBs (Figure 5D, E). The increased expression level of DNA damage marker genes, such as *AtBRCA1, AtNAC103, AtPARP2, AtPARP3*, and *AtRAD17*, in the *fan* mutant further supported our hypothesis that the absence of *FAN* leads to DNA damage (Figure S5).

In mammals, the AATF/Che-1 plays an essential role in the DNA damage response (DDR) pathway (Passananti and Fanciulli, 2007). To test whether *fan* is sensitive to genotoxic agents, we transferred *fan* and Col-0 seedlings (4 DAG) to hydroxyurea (HU, 2 mM), mitomycin C (MMC, 10 µM) and zeocin (20 µM) containing media for 24 hours to induce replication stress, DNA cross-linking and DSBs, respectively. PI-stained roots were imaged with confocal microscopy. Quantification of the area of dead cells in the proximal meristem showed a greater increase in cell death upon HU and MMC treatments in *fan* compared to Col-0 (Figure 5A, F, G), whereas in the cell death response upon zeocin treatment treatment was comparable. Based on these observations we concluded that FAN protects genome integrity and is involved in the DNA damage response upon replication stress and DNA cross-linking.

To confirm this observation, we took advantage of the *CaMV35S* promoter-driven overexpression lines, *FAN*^*OE2*^ and *FAN*^*OE3*^ in which the mRNA level of *FAN*, measured by qRT-PCR, was one hundred times higher than in the control, associated with a longer root meristem length but the same number of root meristem cells compared to Col-0 (Figure S6). Both Col-0 and the overexpression lines accumulated cell death but the extent of the response was weaker in the lines with high *FAN* expression, the number of roots with cell death was lower and the cell death area was also reduced in the *FAN*^*OE2*^ and *FAN*^*OE3*^ compared to Col-0 under HU treatment (Figure S7). Taken together, these results suggest that FAN may help to regulate root growth and maintain genome stability.

Since the *fan* mutant is hypersensitive to genotoxic stress and shows cell death in the root meristem, we asked whether *FAN* gene expression is also affected by genotoxic stress. The transgenic lines *FAN*_*pro*_*:GUS* and *FAN*_*pro*_*:FAN*_*g*_*-GFP* (5 DAG) were transferred to 1/2 MS containing 2 mM HU, 10 µM MMC or 20 µM zeocin and grown for 24 hours. GUS staining indicated that the transcript level of *FAN* did not change significantly upon HU or MMC treatment (1 day), compared to seedlings grown under control conditions (Figure S8A). Alternatively, *FAN* was apparently reduced under 20 µM zeocin treatment due to severe defects in the root meristem (Figure S8A). The GUS staining results were confirmed by qRT-PCR (Figure S8B). Meanwhile, *FAN*_*pro*_*:FAN*_*g*_*-GFP* confocal images showed that HU, MMC, and zeocin reduced FAN protein levels (Figure S8C, S8D).

### FAN partially participates in the ATR-induced DNA damage response

Since the *fan* mutant was hypersensitive to HU treatment, we investigated the effect of replication stress in the *atr-2;fan* double mutant.

The *atr-2* mutant treated with HU (2mM) showed a severe reduction in root growth (Figure 6A-D, S9). In contrast, both root length and meristematic cell number of the *atr-2;fan* were comparable to those of the *fan* mutant, suggesting that the FAN function is essential for the development of the growth inhibition induced by replication stress (Figure 6A-D).

**Figure 6.**
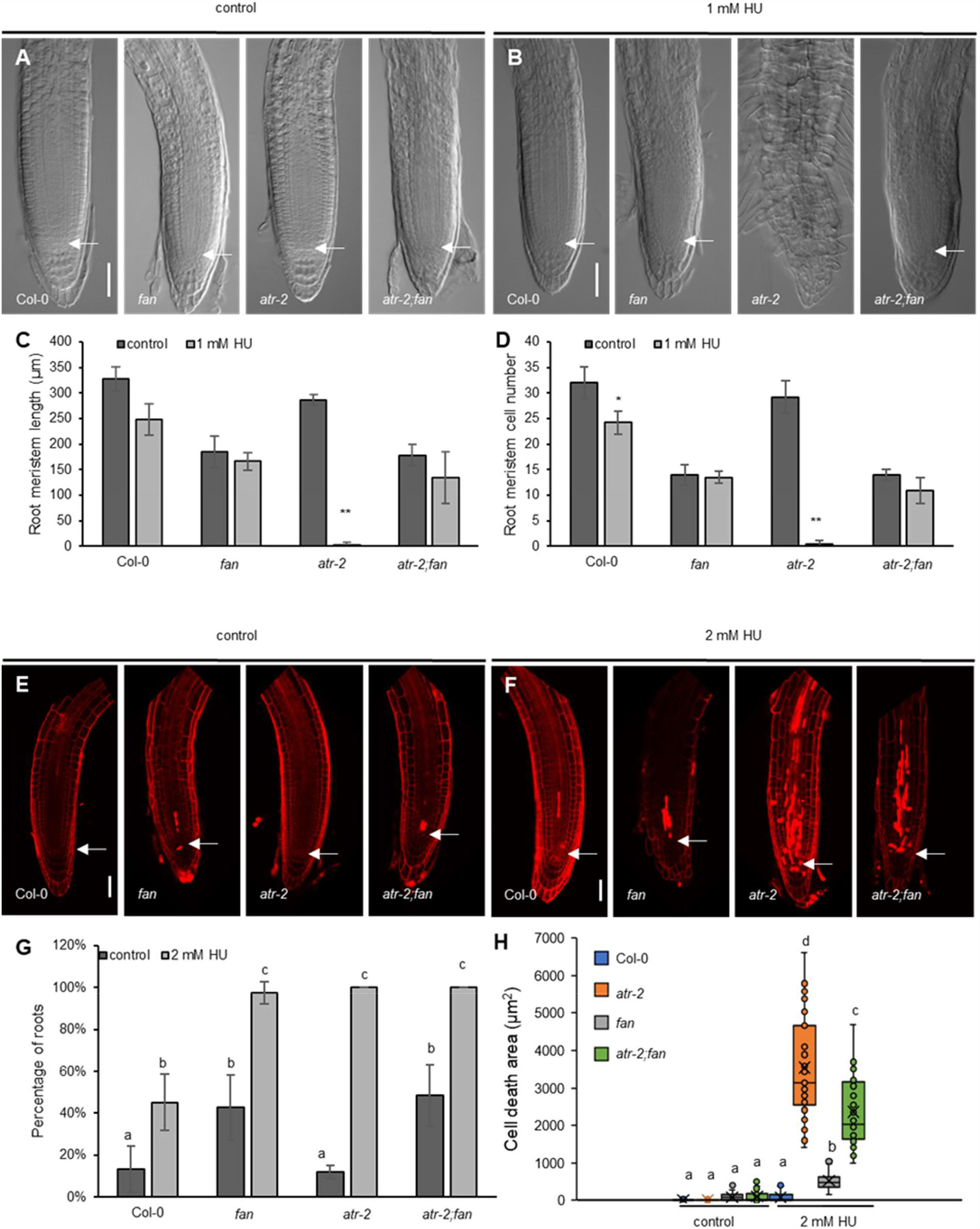
FAN participates in the ATR-induced DNA damage response. (**A**) and (**B**) Representative images of root meristems of 6 DAG Col-0, *fan, atr-2* and *atr-2;fan*. Seedlings grown on 1/2 MS medium with or without 1mM HU. Scale bars = 50 µm; arrowheads point to QC. (**C**) and (**D**) Measurement of root meristem length (**C**) and root meristem cell number (**D**) of 6 DAG Col-0, *fan, atr-2* and *atr-2;fan*. Seedlings grown on 1/2 MS medium with or without 1 mM HU. Data are the mean ± SD of three independent experiments with at least 10 seedlings each. Asterisks indicate significant differences between control and HU treatment (\*\**P* value < 0.01, \**P* value < 0.05, Student’s t test). (**E**) and (**F**) Confocal images of PI-stained root tips of Col-0, *fan, atr-2* and *atr-2;fan* double mutants. 4 DAG seedlings were transferred and grown on 1/2 MS medium with or without 2 mM HU for 24 h. Scale bars = 50 µm; arrowheads point to QC. (**G**) Measurement of the percentage of roots showing dead cells. Data are the mean ± SD of three independent experiments with at least 10 seedlings each. Columns with different letters are significantly different at *P*<0.05 (Duncan’s multiple range means comparisons). (**H**) Measurement of dead cell area from Col-0, *fan, atr-2* and *atr-2;fan* PI-stained roots. The box plots represent cell death are in three independent experiments. The whiskers indicate minimum and maximum values. Crosses indicate mean values. Different letters indicate significant differences at *P*<0.05 (Duncan’s multiple range mean comparisons) between the means of different lines treated with or without HU.

We next analyzed the ratio of roots with /without cell death and measured the cell death area in the *atr-2;fan* double mutant and controls. Although under normal growth conditions, the ratio of roots with cell death was comparable in the *atr-2;fan* and *fan*, and significantly lower in the *atr-2* mutant, (Figure 6E, G, H), upon HU treatment (24h), irrespective of the genetic background, all of the tested seedlings showed a cell death response (Figure 6F, G). In contrast, the area with dead cells differed significantly between the parents and the double mutant. Compared to either to Col-0 or *fan* (Figure 6F), the *atr-2* mutant showed strong cell death response, which, however was greatly reduced in the absence of the FAN function (Figure 6F, G, H). These data suggest that FAN, in partially is required for the ATR-induced DNA damage response to maintain genome integrity in response to replication stress.

## DISCUSSION

AATF/Che-1, an RNA polymerase II binding protein is involved in transcriptional regulation, cell proliferation, DNA damage response, apoptosis and ribosome biogenesis in mammals. Here we show that FAN, the Arabidopsis homolog of AATF/Che-1, is essential for root development and DNA damage response.

### Loss-of-*FAN* maintenance of the SCN

Accumulating evidence indicates that genome integrity is critical for maintaining the stem cell niche. The mutants *teb, fas* and *mms21* that derived from the helicase and DNA polymerase domain encoding gene *TEBICHI*, the *FASCIATA* (*FAS*) gene that encodes subunits of the *Arabidopsis* counterpart of chromatin assembly factor-1 (CAF-1), and the *methyl methanesulfonate sensitivity gene21*, which functions as a subunit of the STRUCTURAL MAINTENANCE OF CHROMOSOMES5/6 complex, all showed severe meristem organization defects associated with cell death response at the root tip (Inagaki et al., 2006; Kaya et al., 2001; Xu et al., 2013). Similar effects were induced by the mutations of *MAIN-related* genes, *MEDIATOR18* and *MERISTEM DISORGANIZATION 1* genes, which encode plant-specific aminotransferase-like plant mobile domain, a subunit of mediator complex, and an essential telomere protein in CTC1 (Cdc13)/STN1/TEN1 complex, respectively (Hashimura and Ueguchi, 2011; Lee et al., 2016; Raya-Gonzalez et al., 2018; Ühlken et al., 2014; Wenig et al., 2013). The RETINOBLASTOMA-RELATED (RBR) is required for both stem cell maintenance and DNA damage response (Biedermann et al., 2017; Horvath et al., 2017; Wildwater et al., 2005).

Here we report, that the mutation in the FAN results in a disorganized SCN and cell death at the root tip, similar to the phenotype developed in *fas*, or *mms21* mutants lacking functional telomeres or chromatin-organizing complex. In addition to aberrations in the SCN, exhaustion of the root meristem, disorganized cell division in the cortex and endodermis cells were observed in *fan*. Induced DNA content in the comet tail, accumulation of γH2AX foci and upregulated DNA damage marker genes indicated genome instability in the *fan* mutant. Similar to the application of DNA damage agents, the absence of FAN function resulted in expressional changes, reducing of expression of essential root development genes, such as *WOX5, SHR, SCR, PLT1* and *PLT2* (Wenig et al., 2013; Xu et al., 2013).

At this stage of the study, it is difficult to conclude whether disturbed expression of root patterning genes, manifested in aberrant root formation, caused DNA damage in the *fan* mutant or whether genome instability leads to DNA damage response, cell death and early differentiation. However, due to the similarity of the *fan* phenotype to that observed in chromatin-organizing protein mutants and the anti-apoptotic function of FAN’s orthologue, AATF/Che-1, in animals, we postulate that FAN has a primary role in determining genome stability and thus root meristem maintenance.

### FAN is involved in the DDR pathway

Based on the fact that AATF/Che-1 is involved in DNA damage response in animals, we investigated whether FAN has a similar role in *Arabidopsis*. Unlike their animal homologs, inactivation of plant genes involved in DDR, in normal growth condition, results in subtle phenotype, but leads to increased sensitivity to genotoxic stress. Similar to *wee1-1* and *atr-2* mutants, upon HU-treatment, *fan* mutant showed elevated level of cell death (Hu et al., 2015).

Since the ATR kinase plays a central role in the response to replication stress (Culligan et al., 2004) we studied the correlation between the FAN and ATR pathway. Upon HU-treatment, compared to *atr-2* mutant, the *atr-2;fan* double mutant not only showed a less severe cell death response, but its root growth was not arrested severely as *atr-2* either. These results indicate that FAN is required to trigger cell death response in the ATR induced pathway upon replication stress.

In mammals, the activity of AATF/Che-1 are post-translationally regulated by phosphorylation, ubiquitination, poly(ADP-ribosyl)ation as well as by conformational changes (Bacalini et al., 2011; Bruno et al., 2006; De Nicola et al., 2007). Further investigations should provide evidence for post-transcriptional and post-translational regulation of FAN in response to DNA damage stress. Similar to its homologue AATF, FAN has a predicted nuclear and a nucleolar localization signal, and is predominantly detected in the nucleolus. Considering the different nucleolar, nucleoplasmic and cytoplasmic functions of AATF, the cellular localization of FAN is an important clue for studying its function. The mouse AATF/Che-1 homologue Traube (Trb) is essential for the growth of preimplantation embryos. The *trb* mutant embryos had a reduced total cell number, arrested development and a lack of ribosomes, polyribosomes and rough endoplasmic reticulum (Thomas et al., 2000). The absence of ribosomes in the *trb* mutant and the nucleolar localization of Traube suggested that Traube is involved in ribosome synthesis. Whether FAN is a transcriptional regulator, a ribosomal synthesis factor or a subunit of a complex will be investigated in the future.

In conclusion, our results show that FAN is involved in the regulation of genome integrity and is essential for stem cell maintenance in *Arabidopsis* (Figure 7).

**Figure 7.**
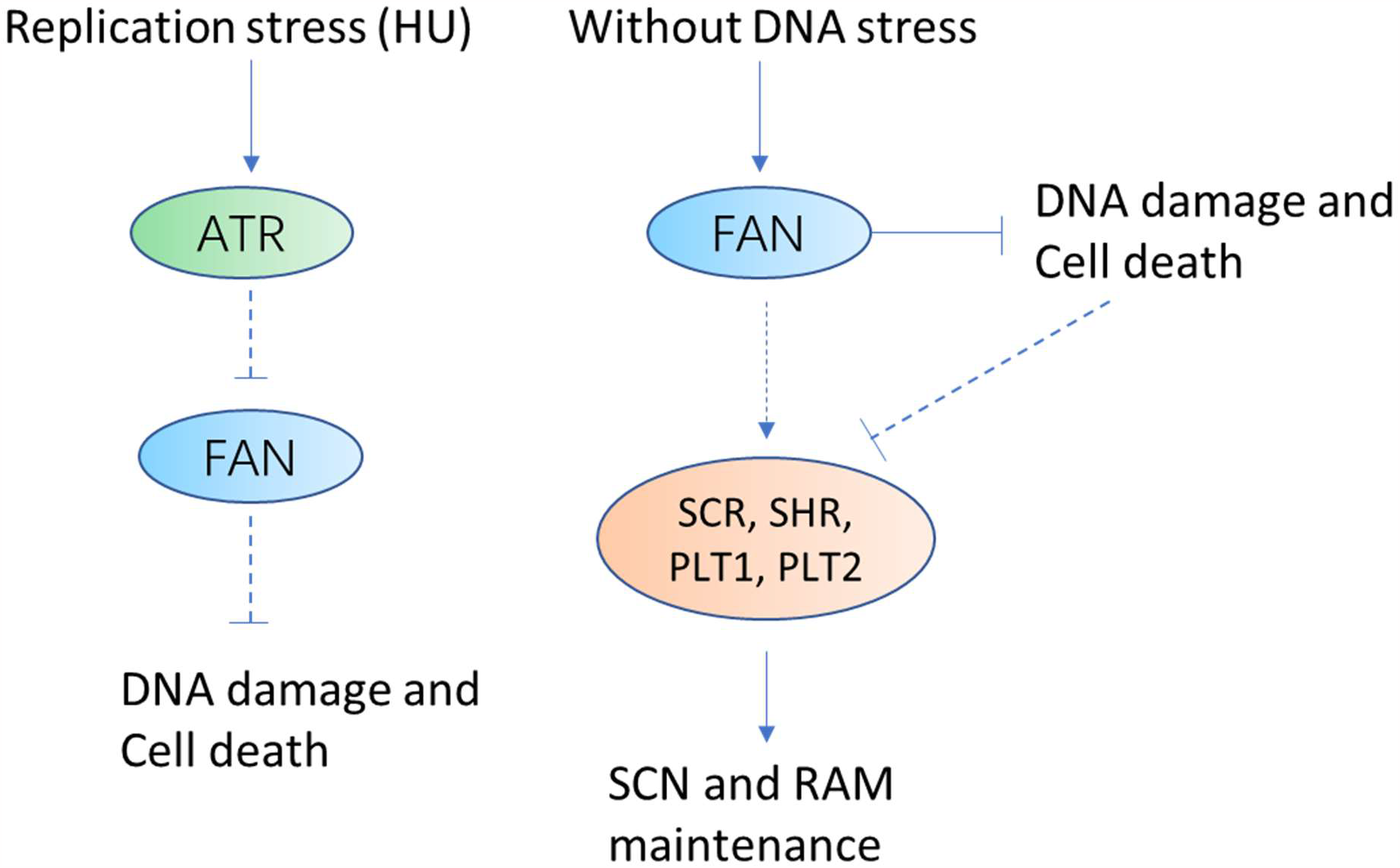
Multiple roles of FAN in root development and DNA damage response. Under non-stress conditions, FAN maintains genome integrity and directly or indirectly promotes the maintenance of SCN and RAM. HU treatment results in the inactivation of FAN and this inactivation may be triggered in an ATR-dependent manner.

## MATERIALS AND METHODS

### Plant materials

The *Arabidopsis thaliana* Columbia-0 (Col-0) ectopy is used as the wild-type. The marker lines used in this work have been described previously. *WOX5*_*pro*_*:GFP* (Blilou et al., 2005), *QC25:GUS* (Sabatini et al., 1999), *SHR*_*pro*_*:SHR-GFP* (Nakajima et al., 2001), *SCR*_*pro*_*:GFP* (Di Laurenzio et al., 1996), *PLT1*_*pro*_*:CFP* (Galinha et al., 2007), *PLT2*_*pro*_*:CFP* (Galinha et al., 2007), *J2341* (Sabatini et al., 2003), *CO2*_*pro*_*:H2B-YFP* (ten Hove et al., 2010), *CO3*_*pro*_*:H2B-YFP* (ten Hove et al., 2010), and *DR5*_*rev*_*:GFP* (Benkova et al., 2003) are ecotype Col-0. The mutants *atr-2* (SALK_032841) and *fan* used in this work are ecotype Col-0. The *35S:H2B-tdTomato* was kindly provided by Prof. Dr. Thomas Laux. The sequence information for the primers used for genotyping is provided in Table S1.

### Protein structure prediction

Nuclear localization signal is predicted by NLS mapper (http://nls-mapper.iab.keio.ac.jp) (Kosugi et al., 2008; Kosugi et al., 2009a; Kosugi et al., 2009b). Nucleolar localization sequence is detected by NOD: NuclOlar localization sequence Detector (dundee.ac.uk) (http://www.compbio.dundee.ac.uk/www-nod/index.jsp) (Scott et al., 2010; Scott et al., 2011). The phosphorylation site of FAN was predicted by the NetPhos 3.1 server (http://www.cbs.dtu.dk/services/NetPhos/) (Blom et al., 1999). Protein domain structure was generated by DOG. Visualization of protein domain structure version 2.0 (Ren et al., 2009).

### EMS mutagenesis and map-based cloning of *FAN*

In this study, the EMS mutagenesis of the *Arabidopsis* population (ecotype: Col-0) and mutant isolation were performed according to (Jander et al., 2003; Yu et al., 2013). To map the *FAN* locus, map-based cloning was performed using standard techniques (Lukowitz et al., 2000). The *fan* mutant (*Arabidopsis thaliana* Col-0 background) was crossed to Landsberg *erecta* to generate the F2 mapping population. Initial coarse mapping localized a region of chromosome 5 between MTE17 and MSN2. Simple sequence length polymorphisms (SSLPs) and derived CAPS (dCAPS) markers were used for fine mapping. Sequence analysis of the genes revealed a G to A mutation in At5g61330 leading to RNA mis-splicing. The wild-type (Col) At5g61330 locus, a 5700 bp Col genomic fragment containing an additional 2197 bp upstream, 2817 bp transcribed region, and 686 bp downstream was introduced into *fan*. We found that this fragment can complement the short root and dwarf seedling of the *fan* mutant. Sequence information of the primers for mapping and mutation site of *fan* are given in Table S2 and Table S3.

### Plasmid construction and generation of transgenic lines

To construct *FAN*_*pro*_*:GUS*, the *FAN* promoter (2197 bp upstream + 201 bp transcribed region) was amplified from BAC MFB13 using primers 164-6 and 164-7. The Pvu I and Asc I cleavage sites were introduced with primers 164-6 and 164-7 respectively. Pvu I is compatible with Pac I. The PCR product was digested by Pvu I and Asc I and *pMDC162* was digested by Pac I and Asc I. The *FAN* promoter fragment was inserted into *pMDC162*. For complementation of mutant plant lines, the *FAN* natural promoter was amplified from BAC MFB13 with full-length genomic DNA and downstream (2197 bp upstream, 2817 bp transcribed region, and 686 bp downstream) using primers 164-35 and 164-11. Sbf I and Sac I cleavage sites were introduced with primers 164-35 and 164-11 respectively. *pCR-Blunt* intermediate vector containing the insertion of the PCR product and *pMDC85* were digested with Sbf I and Sac I. After the digestion reaction, the sequence containing the CaMV35S promoter and the GFP tag between two cleavage sites was removed from *pMDC85*. The fragment was then inserted into *pMDC85* and the *FAN*_*pro*_*:FAN*_*g*_*-FAN*_*3*_*’-UTR* was generated. To analyze the subcellular localization of the protein, the native *FAN* promoter and full-length of genomic DNA without stop codon (2197 bp upstream and 2814 bp transcribed region without stop codon) were amplified from BAC MFB13 using primer pair 164-35 and 164-12. Sbf I and Asc I cleavage sites were introduced with primers 164-35 and 164-12 respectively. *pCR-Blunt* intermediate vector containing the insertion of the PCR product and *pMDC85* were digested with Sbf I and Asc I. After the digestion reaction, the sequence containing the CaMV35S promoter between two cleavage sites was removed from *pMDC85*. The fragment was then inserted into *pMDC85*, resulting in *FAN*_*pro*_*:FAN*_*g*_*-GFP*. For the overexpression line, the CDS (1308 bp without stop codon) was amplified and an additional myc tag was added at the C terminus using primers 164-15 and 164-14. The Spe I and Sac I cleavage sites were introduced with primers 164-15 and 164-

14. The PCR product and *pMDC85* were digested with Spe I and Sac I. After the digestion reaction, the sequence including the GFP tag between two cleavage sites was removed from *pMDC85*. The fragment was cloned into the binary vector *pMDC85* to generate *35S*_*pro*_*:FAN*_*cds*_*-myc*. All vectors were introduced into Agrobacterium strain GV3101 and transformed into *Arabidopsis* via floral dip (Clough and Bent, 1998). Sequence information for the primers used to construct the vectors is provided in Table S4.

### Histochemical analysis

The activity of reporter enzyme β-glucuronidase (GUS) was analyzed in transgenic plants. X-Gluc was used as a substrate for localization of enzyme activity. Seedlings were fixed in 90% acetone for 20 min. Then, seedlings were rinsed with GUS staining buffer (50mg/l X-Gluc; 100 mM NaH_2_PO_4_; 10 mM Na_2_EDTA; 0.5 mM K ferrocyanide; 0.5 mM K ferricyanide; 1‰ Triton X-100 pH = 7.0) and were vacuumed in GUS staining solution for 2 ∼ 3 min. Seedlings were then incubated in GUS staining solution at 37°C. Seedlings were washed twice in PBS buffer and then in 70% Ethanol. Seedlings were mounted in 40% glycerol. Tissues were photographed using an AxioImager A1 microscope (Carl Zeiss Microimaging) and a SteREO Discovery V20 stereomicroscope (Carl Zeiss Microimaging) coupled to an AxionCam MRc digital camera (Carl Zeiss Microimaging).

### Phenotypic analysis

For primary root length, seedlings were scanned and root lengths were measured using Image J (National Institutes of Health; http://rsb.info.nih.gov/ij). For root meristem, roots were mounted in chloral hydrate solution (chloral hydrate: water: glycerol=4:3:1). Root tips were photographed using an AxioImager A1 microscope (Carl Zeiss Microimaging). Meristem length and cell number were measured using Image J, based on the region between the QC and the first elongated cell in the cortex cell file (Casamitjana-Martínez et al., 2003; Dello Ioio et al., 2007). For PI staining of roots, seedlings were rinsed in 10 µg/ml PI for 2 min. Seedlings were then rinsed in water, and roots were mounted in water for analysis by confocal laser scanning microscopy. For Lugol’s staining, seedlings were incubated in Lugol’s solution (1% iodine; 2% potassium iodine) for 5 ∼ 10 min in the dark. Seedlings were then mounted in chloral hydrate solution for observation and imaging (Carl Zeiss Microimaging).

### qRT-PCR

Gene expression was assessed by quantitative RT-PCR (qRT-PCR). Total RNA was extracted from 2 mm root tips of 5 DAG seedlings using the RNeasy Plus Micro Kit (cat. No. 74034 QIAGEN). Prior to cDNA synthesis, the RNA sample was incubated with 1 μl DNase I at 37°C for 30 min (Thermo Scientific), followed by 1 μl 50 mM EDTA at 65°C for 10 min to remove trace amounts of DNA. The amount of RNA was measured using a PRQlab NanoDrop 1000 spectrophotometer. The integrity of RNA was checked by agarose gel electrophoresis. Reverse transcription was performed using Thermo Scientific ReverAid Reverse Transcriptase (EP0041). qRT-PCR was performed using Maxima SYBR Green qPCR Mastermix (Thermo Scientific K0222) on an ABI 7300 real-time PCR system (Applied Biosystems). Primers used for qRT-PCR are listed in Table S5. *UBIQUITIN10* was used as a reference gene.

### Confocal microscopy

GFP, CFP, Alexa Fluor 488 picolyl azide, DAPI, tdTomato, and PI fluorescence were photographed using a Zeiss LSM 510 microscope (Carl Zeiss Microimaging) and an A1 HD25/A1R HD25 confocal microscope (Nikon). 488nm line laser was used for excitation and emission was detected at 505 to 530 nm for GFP and Alexa Fluor 488 picolyl azide. 458nm line laser was used for excitation and emission was detected at 465 to 520 nm for CFP. For PI and tdTomato, the lase line 561 nm and 590 to 653nm were used for excitation and emission respectively. For DAPI, the excitation wavelength is 408 nm and the emission wavelength is 480 nm. Images were processed and exposed using ZEN lite (Carl Zeiss Microimaging) and NIS-Elements Viewer (Nikon). Fluorescence intensity and plot profile were measured using Image J software (National Institutes of Health; http://rsb.info.nih.gov/ij) and Fiji (Schindelin et al., 2012).

### Comet assay

The method for comet assay was established according to (Menke et al., 2001), (Olive and Banath, 2006), and (Yoshiyama et al., 2017). Briefly, 7-day-old seedlings were cut with a fresh razor blade on ice in PBS buffer containing 50 mM EDTA. Nuclear suspensions were filtered through a 40 µm strainer. The nuclear suspension was mixed with liquid 1% low melting point agarose at a ratio of 1:10 (v/v). Two drops of 50 µl of the resulting nuclear suspension were applied separately to the precoated slide and covered with 22 mm × 22 mm coverslips. The slides were then incubated at 4°C for 60 min. Slides were incubated in neutral lysis solution (high salt: 2.5 M NaCl, 10 mM Tris-HCl, pH 7.5, 100 mM EDTA) for 20 min (N/N protocol) at room temperature. This was followed by equilibration in 1×TBE buffer for 5 min and electrophoresis at room temperature in the same buffer for 6 min at 25 V (1 V/cm). Excess electrophoresis solution was drained from the slides and gently immersed twice in dH_2_O for 5 min each followed by70% ethanol for 5 min. Slides were dried on the sterile bench. Dry agarose gels were stained with 10 µg/ml of PI and covered with coverslips. Comets were observed using a fluorescence microscope-AxioImager A1 (ZEISS). Comet analysis was performed using the TriTek Comet Score software. We measured at least 100 comet images from each slide and avoided analyzing doublets or comets at the edge of the slide.

### Immunofluorescence staining

5 DAG plants were incubated for 30 min in fixation buffer (4% paraformaldehyde in 100 ml MTSB) in the cold room (4?). Cell wall digestion, blocking, primary antibody incubation, secondary antibody incubation, and nuclear co-staining of the nucleus were performed as previously described (Pasternak et al., 2015) and (Biedermann et al., 2017). A rabbit anti-plant γH2AX antibody (Amiard et al., 2010) was diluted to a working concentration of 1:500. A Goat anti-rabbit IgG (H+L) highly cross-adsorbed secondary antibody, Alexa Fluor 488 (Invitrogen, A-11034), was used at a 1:500 dilution.

### Accession numbers

Sequence data from this article is available from the *Arabidopsis* Genome initiative under the following accession numbers: At5g61330 (*FAN*), At3g11260 (*WOX5*), At4g05320 (*UBQ10*), At3g20840 (*PLT1*), Atg51190 (*PLT2*), At3g54220 (*SCR*), At4g37650 (*SHR*), At5g40820 (*ATR*), At4g21070 (*BRCA1*), At4g02390 (*PARP2*), AT5G22470 (*PARP3*), AT5G64060 (*NAC103*), and AT5G66130 (*RAD17*).

## ACKNOWLEDGEMENTS

We thank Prof. Dr. Yan Zhang for comments on the manuscript. This work was supported by initiation fund from the Shandong Agricultural University; National Natural Science Foundation of China (31570291); Shandong “Foreign experts double hundred” Program (WST2017008) and Natural Science Foundation of Shandong Province (ZR2021MC175). In addition the work was supported by Bundesministerium für Bildung und Forschung [BMBF Haploswitch, Innobeet]; the Excellence Initiative of the German Federal and State Governments [EXC 294, SFB746]; and the Deutsches Zentrum für Luft und Raumfahrt [DLR 50WB1022]. Beatrix. M. Horvath was funded by a Marie-Curie IEF fellowship (FP7-PEOPLE-2012-IEF-330789).

## AUTHOR CONTRIBUTIONS

X.L. and K.P. designed the research and supervised the experiments; X.L. and F.L. wrote the manuscript; B.M.H improved the preliminary version of the manuscript; F.L., B.W and X.W. performed most of the experiments; B.K.H, B.M.H., L.D.Y., D.D. and S.Q. contributed reagents, materials, analytical tools and suggestions.

## FIGURE LEGENDS

**Figure S1. Abnormal shape of the QC in the *fan* mutant**.

DIC images of Lugol stained roots showing the expression of *QC25:GUS* in Col-0 and *fan* (6 DAG). Scale bar = 50 µm, arrowheads point to QC.

**Figure S2. *FAN* genomic DNA fragment can complement the *fan* phenotype**.

(**A**) Complementation of the *fan* mutation with the *FAN* genomic DNA fragment expressed under its own promoter. Root growth phenotype of Col-0, *fan* and the complementation lines FAN C1 and FAN C2 (6 DAG). Scale bar = 1 cm. (**B**) Primary root length. (**C**) Root meristem of Col-0, *fan* and complementation lines. Scale bar = 50 µm; arrowheads point to QC and the first elongated cortical cell. (**D**) amplifying cortex cell number and (**E**) root meristem length. Data represent the mean ± SD, (n>20). Columns with different letters are significantly different at *P*<0.05 (Duncan’s multiple range means comparisons).

**Figure S3. FAN is the conserved AATF/Che-1 in *Arabidopsis***.

Evolutionary analysis was performed in MEGA X and the history was inferred using the neighbor-joining method. The percentage of replicate trees in which the associated taxa clustered together in the bootstrap test (500 replicates) are shown next to the branches.

**Figure S4. Expression levels of *PLT1, PLT2, SHR* and *SCR* are reduced in *fan* compared to Col-0**.

(**A**) and (**B**) Mean grey value of *PLT1*_*pro*_*:CFP* and *PLT2*_*pro*_*:CFP* in Col-0 and *fan*. Data are the mean ± SD, asterisks indicate significant difference compared to Col-0. (\*\**P* value < 0.01, Student`s t-test, n>10). (**C**) and (**D**) Mean grey value of *SHR*_*pro*_*:SHR-GFP* and *SCR*_*pro*_*:GFP* in Col-0 and *fan*. Data are the mean ± SD, asterisks indicate significant differences compared to Col-0. (\*\**P* value < 0.01, Student’s t test, n>10).

**Figure S5. The increased expression level of DNA damage marker genes in *fan* mutant compared to Col-0**.

The relative expression level of DNA damage marker genes in *fan* compared to Col-0 (5 DAG) measured by qRT-PCR. 2 mm root tips were collected as samples. Data are the mean ± SD of three independent biological replicates.

**Figure S6. Root meristem phenotype of *FAN* overexpression lines**.

(**A**) Representative images of the root meristem of Col-0, *FAN* overexpression lines *FAN*^*OE2*^, and *FAN*^*OE3*^ (6 DAG). Scale bar = 50 µm; arrowheads point to QC and the first elongated cortex cell. (**B**) Relative mRNA levels of *FAN* in Col-0, *FAN*^*OE2*^ and *FAN*^*OE3*^ (5 DAG) grown on 1/2 MS medium were analyzed by qRT-PCR using three independent biological replicates. (**C**) Quantification of root meristem length of Col-0, *FAN*^*OE2*^ and *FAN*^*OE3*^ (6 DAG). The box plots represent the root meristem length. Whiskers indicate minimum and maximum values. Crosses indicate the mean values. Asterisks indicate significant difference compared to Col-0 (\*\**P* value < 0.01 Student’s t test, n>10). (**D**) Quantification of number of root meristem cells of Col-0, *FAN*^*OE2*^, and *FAN*^*OE3*^ (6 DAG). Data are the mean ± SD, n>10)

**Figure S7. Decreased cell death level of *FAN* overexpression lines under HU treatment**.

(**A**) and (**B**) PI-stained root tips of Col-0, *FAN*^*OE2*^ and *FAN*^*OE3*^ under control conditions (**A**) and 2 mM HU treatment (**B**). 4 DAG seedlings were transferred to 1/2 MS medium with or without 2 mM HU and grown for 1 day. Scale bars = 50 µm; arrowheads point to QC. (**C**) Ratio of roots (%) with cell death response, graph shows means ± SD from three independent experiments with at least 10 seedlings each. Asterisks indicate significant differences compared to control, respectively (\*\**P* value < 0.01, \**P* value < 0.05, Student’s t test). (**D**) Quantification of dead cell area (µm^2^) from PI-stained roots of Col-0 and *FAN* overexpression lines. The box plots represent cell death are in three independent experiments. The whiskers indicate minimum and maximum values. The crosses indicate mean values.

**Figure S8. *FAN* transcript level is not effected by DNA damage reagent treatment**.

(**A**) GUS-stained images of *FAN* promoter fused to *GUS* reporter gene (*FAN*_*pro*_*:GUS*) seedlings. 5 DAG seedlings were transferred and grown for 1 day on 1/2 MS medium with either no stress, 2 mM HU, 10 µM MMC, or 20 µM zeocin respectively. Scale bar = 50 µm. (**B**) Relative mRNA levels of *FAN* under DNA damage stress treatments were measured by qRT-PCR. 4 DAG seedlings were transferred to 1/2 MS liquid medium without stress, 2 mM HU, 10 µM MMC, or 20 µM zeocin and grown for 24 h. After that, 5 DAG seedlings were collected as samples. Data are the mean ± SD of three independent biological replicas. Columns with different letters are significantly different at *P*<0.05 (Duncan’s multiple range means comparisons). (**C**) Confocal images of PI-stained of *FAN*_*pro*_*:FAN*_*g*_*-GFP* root tips. 5 DAG seedlings were transferred to 1/2 MS plates with no stress, 2 mM HU, 10 µM MMC or 20 µM zeocin and grown for 24 h. Magenta, PI staining and green, GFP. Scale bar = 50 µm; arrowheads point to QC. (**D**) Mean grey value of *FAN*_*pro*_*:FAN*_*g*_*-GFP* under DNA damage treatment. Data are the mean ± SD, n>10. Columns with different letters are significantly different at *P*<0.05 (Duncan’s multiple range means comparisons).

**Figure S9. Primary root length of *atr-2;fan* double mutant is partially rescued compared to *atr-2* mutant upon HU treatment**.

(**A**) to (**D**) Primary root phenotype of 6 DAG Col-0 (**A**), *fan* (**B**), *atr-2* (**C**) and *atr-2;fan* (**D**) seedlings grown on 1/2 MS medium containing 0 (control condition) or 1 mM HU. Scale bars = 1 cm. (**E**) Measurement of primary root length of Col-0, *fan, atr-2* and *atr-2;fan* (6 DAG). Data are the mean ± SD from three independent experiments with at least 10 seedlings each. Asterisks indicate significant differences compared to the control. (\*\**P* value < 0.01, \**P* value < 0.05, Student’s t test).

**Table S1. Primers for genotyping**.

**Table S2. The sequence information for the primers used for mapping**.

**Table S3. The sequence information for the primers used for mapping**.

**Table S4. Primers for vector construction**.

**Table S5. Primers for qRT-PCR**.

