## supplemental figures and tables for "FAN, the homolog of mammalian Apoptosis Antagonizing Transcription Factor AATF/Che-1 protein, is involved in safeguarding genome stability through the ATR induced pathway in *Arabidopsis*"

**Table S1. Primers for genotyping**

| Primer names | Sequence (from 5‘ to 3‘) |
|---|---|
| For *atr2*: (SALK_032841) | |
| SALK_032841_LP | GCAGCAAAAATTTCTTGGTTG |
| SALK_032841_RP | ACTTCAAGGGTTCCGATGTTC |
| For *fan* | |
| fan mutant F | TAAGCTATGCTCTTCTCCTCTTC |
| fan mutant R | CCTCTGGATTTGGTTCCAT |

**Table S2. The sequence information for the primers used for mapping**

| Markers | Name | Sequence from 5' to 3' |
|---|---|---|
| MSN2 | 457122_F1: | TAAATGGGCGAGACAACATCACAG |
| | 457122_F2: | CTATAGTGGGACTGAAAAGCCC |
| | 457122_R1: | GGAAGTTGGTGAAAGAGCGGC |
| | 457122_R2: | GTAGCAAACTCATCGCGTTGGGG |
| MUB3 | 457274_F1: | GGGCATAGAATAAGAGCATATTCC |
| | 457274_F2: | GAATAAGGCAGTGGCAACCATTG |
| | 457274_R1: | GAGAATACAATATTCGTGAGAG |
| | 457274_R2: | CCATCACTCAAATTTGAACGTTGC |
| K11J9 | 454021_F1: | CGTTTTACGAGCATGGTCTTGGC |
| | 454021_F2: | CATGGTCTTGGCTAACATCCTCC |
| | 454021_R1: | CCAAACTCCTCGTGTTTGGCTG |
| | 454021_R2: | TGGCTGACCGGTTATGATGAGg |
| F15L12 | 449419_F1: | CCTGACTATCCTCACTTCTGAGG |
| | 449419_F2: | TTGAGGTTCATGTTTCTATGACG |
| | 449419_R1: | GCCAAAGGGTAAGTGGGTGGGTGA |
| | 449419_R2: | CAGAGACAGAGACGGTGAGTTAGAC |
| MTE17 | 457148_F1: | GGGTTCACGTGAACATGACAC |
| | 457148_F2: | GATCGATCGATAGGAAGTGTTGG |
| | 457148_R1: | CTGAAGCTGTGTTGCTCGTTGAG |
| | 457148_R2: | GCGGGGTGATGGAGAGATTCAC |
| MTH12 | MTH12_F: | CGGCATCTGTTCATGCATTATA |
| | MTH12_R: | CTCAAGGCTAAGTAGTGTATGA |

| | | |
|---|---|---|
| MMN10 | MMN10_F: | CCCCATTGCCCCGCGGTAATAAGC |
| | MMN10_R: | CGAACCATCACCACTGGTGAGTG |
| F15L12 | F15L12_F: | AGTAGGAATTGGAATGAGCAA |
| | F15L12_R: | TGTCATGTGAGGACAAGTCTGAA |
| MUP24 | MUP24_F: | GTAGAATCAGAAATACCATAATC |
| | MUP24_R: | GGTGTCCAATCAAGTTTTCGGTT |
| MAE1 | MAE1_F: | GAACTGGACTATAATATAAATT |
| | MAE1_R: | GACTTGGGTCAGAGGACAAACG |
| MSL3 | MSL3_F: | CTTACGTTTGTGAACACATATTA |
| | MSL3_R: | GACAAAATTGCAAACGCGGAGAT |
| MFB13 | MFB13_F: | GACGACTGATTACATAACATAGT |
| | MFB13_R1: | GTGTGTATTGTCTATATATATTACG |
| | MFB13_R2: | AGCTGTCATGCGTGATGCTTG |
| MAC9 | MAC9_F: | CACATGAGACTTGAGTGTTGTTC |
| | MAC9_R: | CGGATTTCATACGAACTTGCTAA |
| K19B1 | K19B1_F: | GGTTATCGAATATATAAAATGTG |
| | K19B1_R: | GTCAATGTTGGGAATTTGCAGTCC |
| MQB2 | MQB2_F: | GGCGACTACTAGCATAAAAAATA |
| | MQB2_R: | GATCTTGCCATTTATTTGGTCAA |
| MBK5 | MBK5_F: | GGCCCATCTAGAGTATAACCATG |
| | MBK5_R: | CAGCTACTGCGTGCAAATAAAGAT |
| MGI19 | MGI19_F: | CTAGAGAGACAAGATAAGACACC |
| | MGI19_R: | GTTATCGCCAAACTTGACCCTTA |

| | | |
|---|---|---|
| MUB3 | MUB3_F: | GCCTGGCTGATATTATGAACTTTC |
| | MUB3_R: | GGCTATTATCACTTCCGAAGAGGTT |
| MUB3 | 457274_F3: | GCATTGAATAAGGCAGTGGCAACC |
| | 457274_R3: | CGTGTGGCAAATCCAATAGTTAG |
| MAF19 | MAF19_a_F: | CTCAACCAATCAAAGGCGGACACC |
| | MAF19_a_R: | TATTAAACGATAAATTCGCCGTTTGC |
| MAF19 | MAF19_b_F: | GTCACTGTTGTCTTTCTAGAAACAGAG |
| | MAF19_b_R: | CTCTATCTCTCTCTGTGTCTCTCCA |
| 10A10 | 10A10_F: | GGTCACAGGGATCAAGATGTGG |
| | 10A10_R: | GCCTTATGGATTTTCTGGAGAAAG |
| EG7F2 | EG7F2_F: | GCATAGAATTTGACGATAACGAGC |
| | EG7F2_R: | GATCTGTGTAGGACTACGAGAC |

**Table S3. The sequence information for the primers used for mapping**

| Name | Sequence from 5' to 3' |
|---|---|
| 210_F1: | GGGTCGTTCTGAGTCGTCTCC |
| 210_R1: | GTCTCTCAGGATAATATCAC |
| 210_F2: | GTAGTAACCAATCCAAGTGTTCC |
| 210_R2: | CTCACACCAACTAATGTCCTCAG |
| 210_F3: | GCCGTTGAAATCGACCATGATC |
| 210_R3: | CTCCACTCTCTGATAAGCATC |
| 210_F4: | GGGACTGAACTCAGCACCCAGAG |
| 210_R4: | ACTGGAAAGATTCTTATCATG |
| 200_F1: | GGTAATGGAGAGATTCTCATG |
| 200_R1: | CTGGTTTATACAGCAGGAAGAGGAAC |
| 190_F1: | TCGACGAGATAGTGAGAGGAGTA |
| 190_R1: | ATTCCCCTGTGGCTCCACTGC |
| 190_F2: | CCAGAGAAAAGCATGTTCCAAGAGG |
| 190_R2: | CACTTGCATTACGCAGTGCAGTCA |
| 190_F3: | GATATTATAGCTTTGGATTAGGTAC |
| 190_R3: | GACAAAGTTTAGATGCAGCTGGTC |
| 190_F4: | GGAGAAACTAAGCATTCATGGGAGG |
| 190_R4: | GATCGGTTTCTTCTCGGATTATCCTTG |
| 190_F5: | CGAGGGGCACCAGAAGATAAAGTGG |
| 190_R5: | CATCCATTGTTGACAACTTAAACATG |

| | |
|---|---|
| 180_F1: | ACAATCTCTAAACCCTGACCTCCC |
| 180_R1: | CGATTAGGGGGTCCTGCAGATGCGC |
| 170_F1: | AGTATCGCCGACGCCGCAGCACA |
| 170_R1: | GGGAATTCTACTACTGGTTTTCATC |
| 160_F1: | TTAACTCCATCTGATACTCAGCTG |
| 160_R1: | GGGTCACCACTACGTGCCTTCACCATG |
| 160_F2: | CACGCGCTGAGCTTCACTTGTCAA |
| 160_R2: | AGGGGGAAATAGATTCACTGATCATG |
| 150_F1: | CGTCGGAGCTCAGATCACGAGCCG |
| 150_F1: | GTTCATCCTGCTCAACGTCATTCCGAAC |
| 150_F2: | GAGGTTCAAGTTGCACAGTCAGACG |
| 150_R2: | GCATCAGAAGTACTAGTAATAGGC |
| 150_F3: | ACATATGTAAGTGAACACTTCTTC |
| 150_R3: | CATCAAGAGCATCTTCCAAGTAGCC |
| 150_F4: | ACAAAGCTGAGTCAGGCGAGGGAG |
| 150_R4: | AACGAGTCTAATTCCAGACTACAT |

**Table S4. Primers for vector construction**

| Primer names | Sequence (from 5‘ to 3‘) |
|---|---|
| Promoter reporter analysis with GUS | |
| Forw: 164-6: | GGGGCGATCGGCGGCCGCGGAGACAGAGATATCATATCCATAGCTC |
| Rev: 164-7: | GGGGGGCGCGCCAAATCAGGTAATTGATCATCTTCATTGTCGCT |
| Complementation analysis of mutant plant lines | |
| Forw: 164-35: | GGGGGCCTGCAGGGGAGACAGAGATATCATATCCATAGCTC |
| Rev: 164-11: | GGGGGAGCTCGTGAAAGAGACGCAGAAGATGTGAAACC |
| Subcellular localization | |
| Forw: 164-35: | GGGGGCCTGCAGGGGAGACAGAGATATCATATCCATAGCTC |
| Rev: 164-12: | GGGGGGCGCGCCAAGCTTCAGACTGAACGTTTCTGGTCTTG |
| Overexpression of CDS | |
| Forw: 164-15: | GGGGACTAGTATGGCTGGGGGGTCAAAGAGGTCT |
| Rev: 164-14: | GGGGGAGCTCTTAAAGATCCTCCTCAGAAATCAACTTTTGCTC AGCTTCAGACTGAACGTTTCTGGTCTTG |

**Table S5. Primers for qRT-PCR**

| Gene names | Forward Primers (from 5' to 3') | Revers Primers (from 5' to 3') |
|---|---|---|
| *PLT1* | TTGCAGCAACAGTCGAGCCAGA | TCGGTCGATCCAACAGTCGAGC |
| *PLT2* | GGCTGAGGAAGAGTTTCCAGCCG | TCCATACCCTACCTTGCCTCGC |
| *RAD17* | GCGCCCGTTGTTCTTTTGAT | GTCGTGCAGTCTGGTCAGAA |
| *BRCA1* | TGAAGCTGATGGGAAGCAGG | GTTTCTCGGTGGGACTTCGT |
| *NAC103* | AAACAAGGAAGCTCCGGTGT | GAGCTGGAGGAGAAGGAGGA |
| *SCR* | AGCAACAACCGTGGTCCTCCT | ATACAGTAGCCGCCGCCGTAA |
| *SHR* | GCGAACGATGCTACCGAACCAT | GCCTCTCCGTCTACTGCTTCCA |
| *PARP3* | GCTTTTGAGACGGTGAGGGA | GCACTGACACACCGTACTCA |
| *PARP2* | TCCTGAAGCGCCTGTAACTG | TGCTGTTTTCCCCACACCTT |
| *UBQ10* | CCTGCGTCTTCGTGGTGGTT | GTCGAGTCACTTTGCAGGCGT |

**Figure S1. Abnormal shape of the QC in the *fan* mutant**.

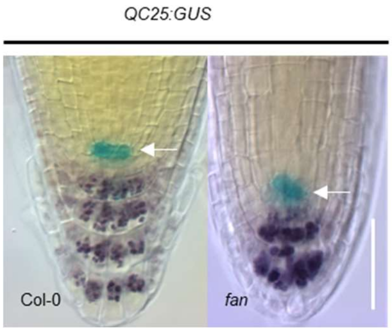

**Figure S2.** ***FAN*** **genomic DNA fragment can complement the** ***fan*** **phenotype**.

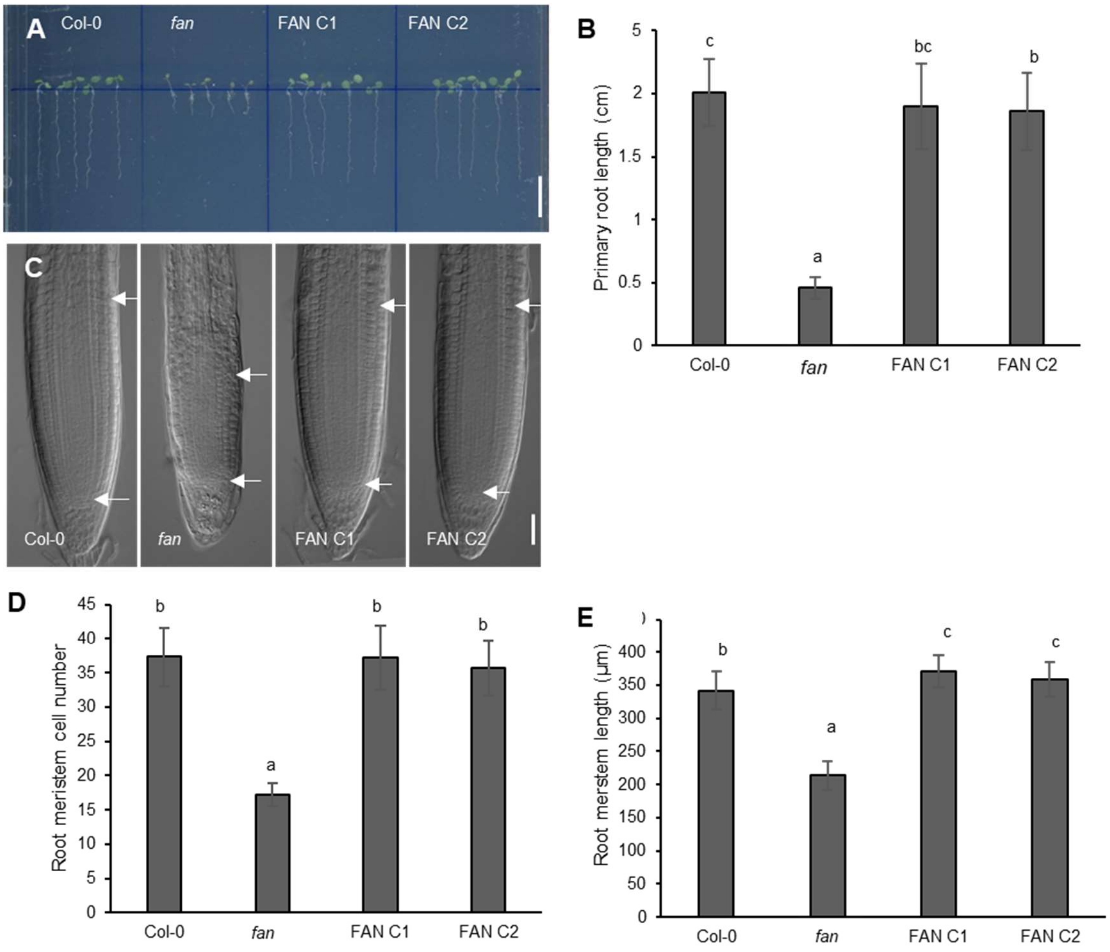

**Figure S3. FAN is the conserved AATF/Che-1 in *Arabidopsis*.**

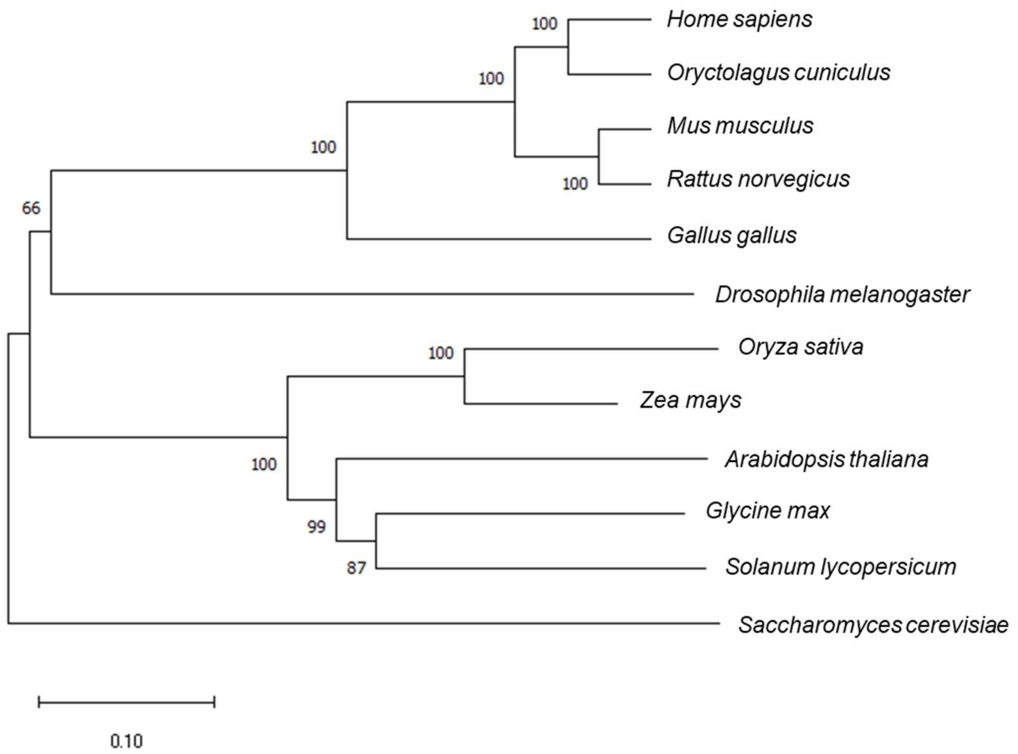

**Figure S4. Expression levels of *PLT1*, *PLT2*, *SHR* and *SCR* are reduced in *fan* compared to Col-0.**

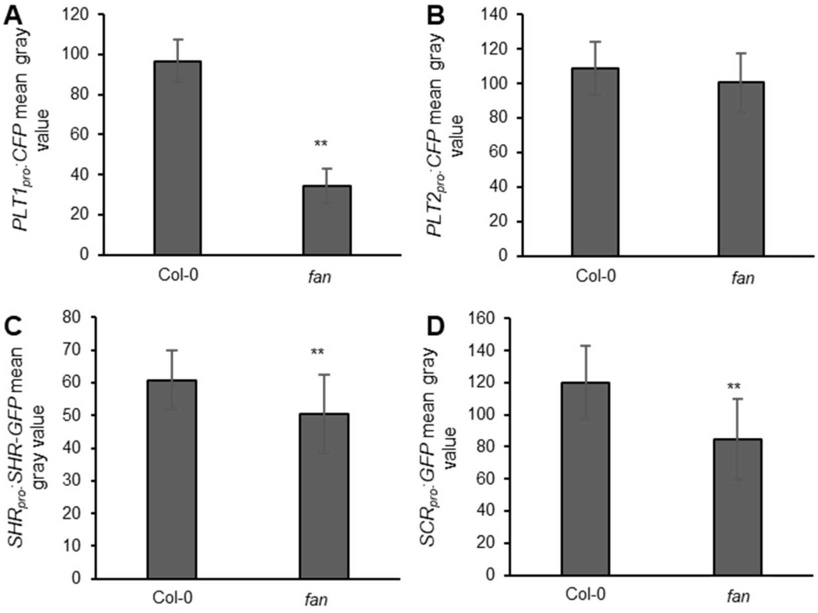

**Figure S5. The increased expression level of DNA damage marker genes in *fan* mutant compared to Col-0.**

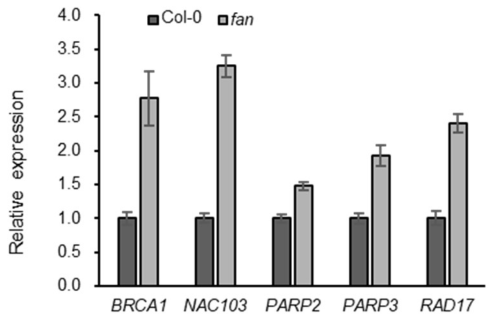

**Figure S6. Root meristem phenotype of *FAN* overexpression lines.**

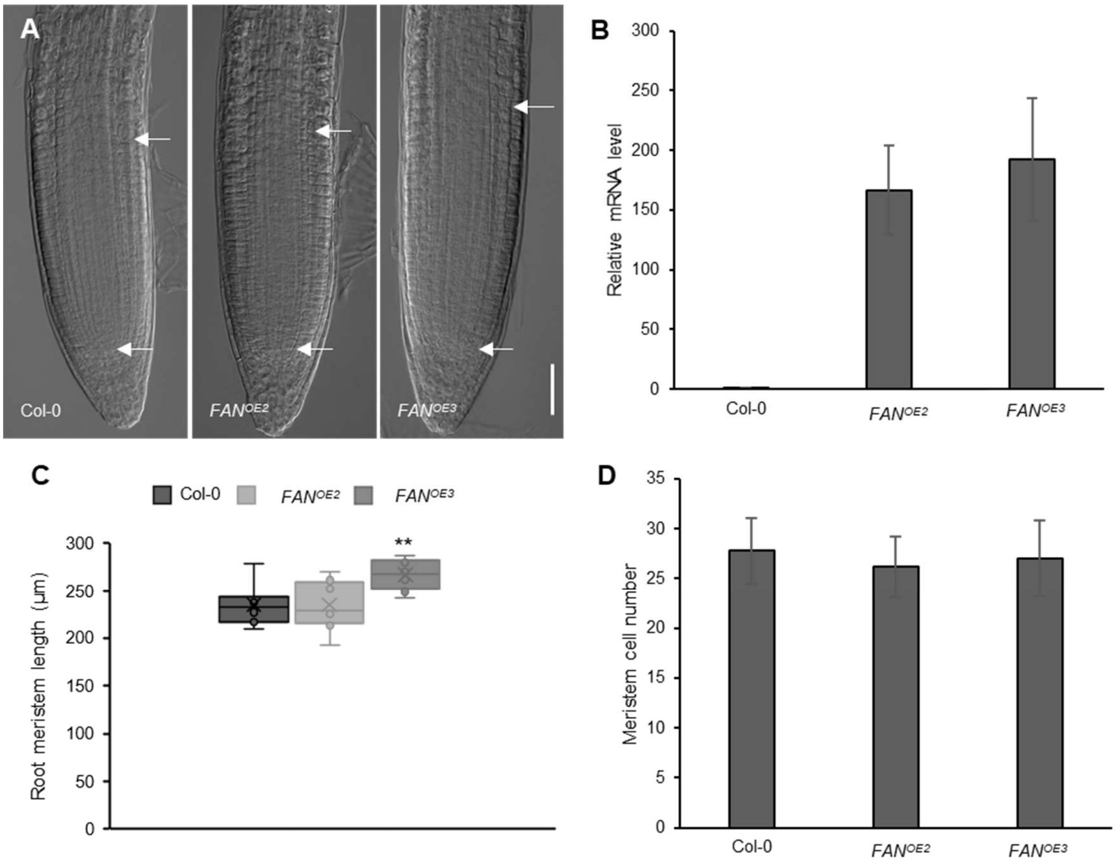

**Figure S7. Decreased cell death level of *FAN* overexpression lines under HU treatment.**

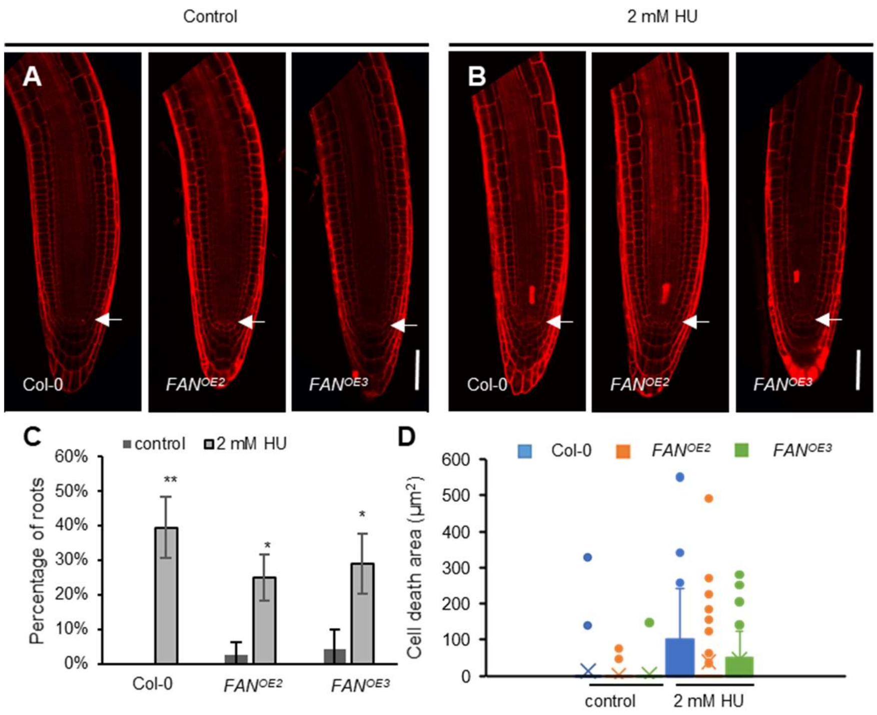

**Figure S8. *FAN* transcript level is not effected by DNA damage reagent treatment.**

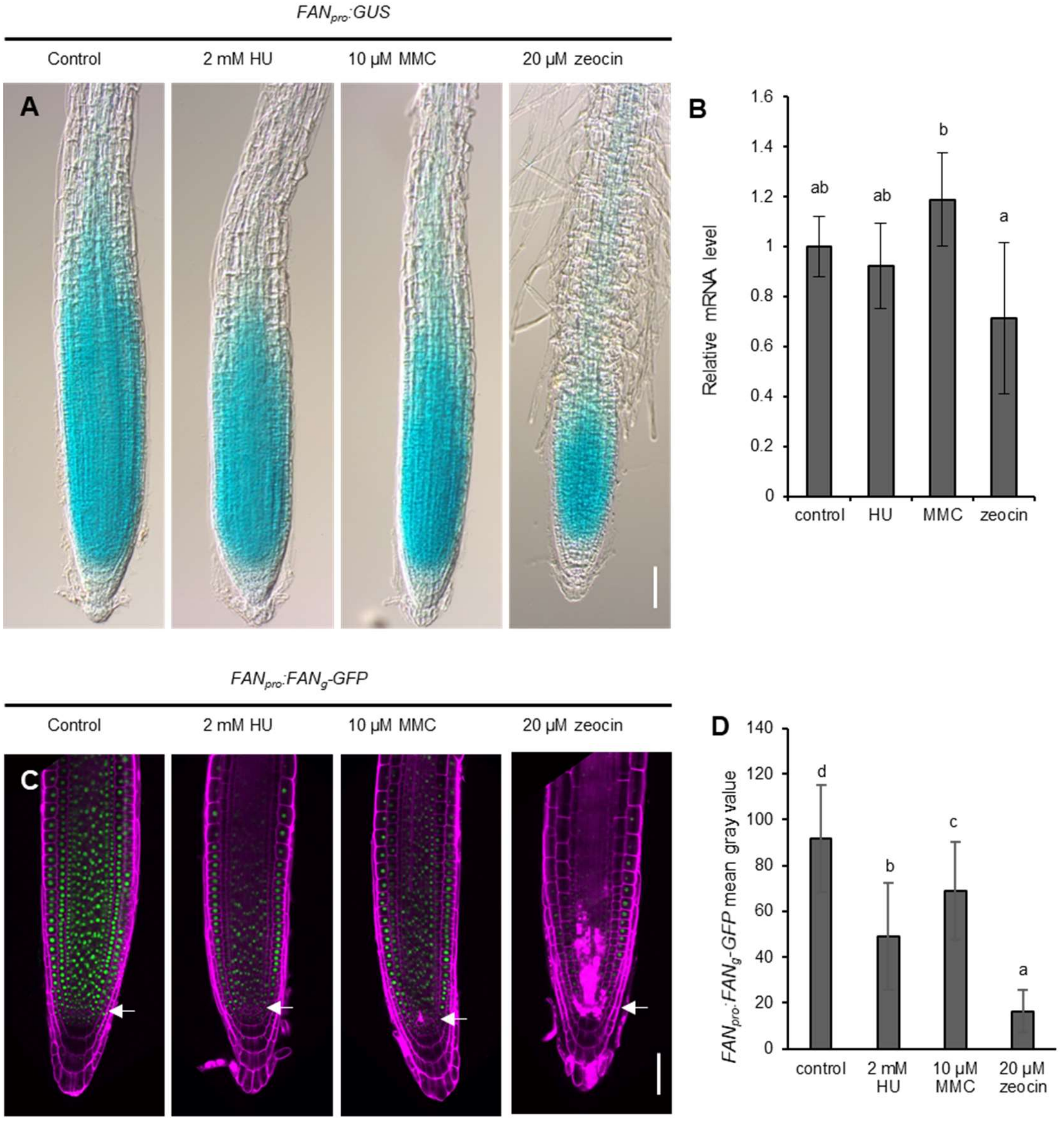

**Figure S9. Primary root length of *atr-2;fan* double mutant is partly rescued compared to *atr-2* mutant upon HU treatment.**

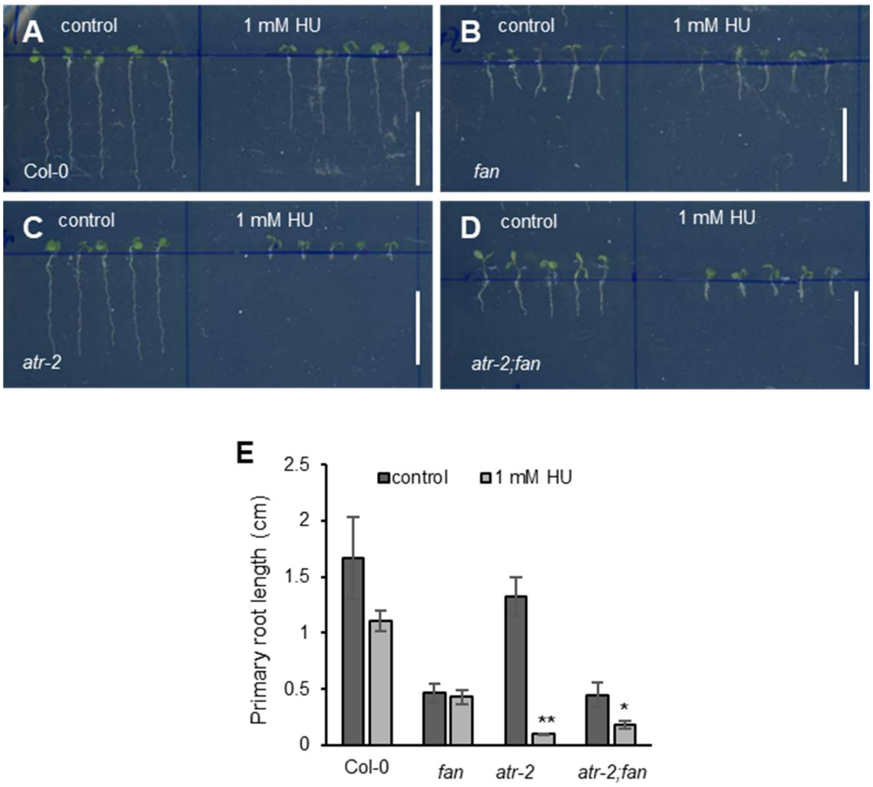
